# A simplified cell-based assay to identify coronavirus 3CL protease inhibitors

**DOI:** 10.1101/2020.08.29.272864

**Authors:** Samuel J. Resnick, Sho Iketani, Seo Jung Hong, Arie Zask, Hengrui Liu, Sungsoo Kim, Schuyler Melore, Manoj S. Nair, Yaoxing Huang, Nicholas E.S. Tay, Tomislav Rovis, Hee Won Yang, Brent R. Stockwell, David D. Ho, Alejandro Chavez

**Affiliations:** Department of Pathology and Cell Biology, Columbia University Irving Medical Center, New York, NY, 10032, USA; Medical Scientist Training Program, Columbia University Irving Medical Center, New York, NY, 10032, USA; Aaron Diamond AIDS Research Center, Columbia University Irving Medical Center, New York, NY, 10032, USA; Department of Microbiology and Immunology, Columbia University Irving Medical Center, New York, NY, 10032, USA; Department of Biological Sciences, Columbia University, New York, NY, 10027, USA; Department of Chemistry, Columbia University, New York, NY, 10027, USA

## Abstract

We describe a mammalian cell-based assay capable of identifying coronavirus 3CL protease (3CLpro) inhibitors without requiring the use of live virus. By enabling the facile testing of compounds across a range of coronavirus 3CLpro enzymes, including the one from SARS-CoV-2, we are able to quickly identify compounds with broad or narrow spectra of activity. We further demonstrate the utility of our approach by performing a curated compound screen along with structure-activity profiling of a series of small molecules to identify compounds with antiviral activity. Throughout these studies, we observed concordance between data emerging from this assay and from live virus assays. By democratizing the testing of 3CL inhibitors to enable screening in the majority of laboratories rather than the few with extensive biosafety infrastructure, we hope to expedite the search for coronavirus 3CL protease inhibitors, to address the current epidemic and future ones that will inevitably arise.

## Introduction

The outbreak of a novel coronavirus (SARS-CoV-2) infection of the past several months has paralyzed countries around the world^1,2^. This crisis is further exacerbated by the dearth of approved therapeutics, leaving physicians with few proven treatment options. In the past two decades, the world has already suffered from two other major coronavirus outbreaks, Severe Acute Respiratory Syndrome (SARS) and Middle East Respiratory Syndrome (MERS), suggesting that coronaviruses represent a real and ever-present threat to global health that must be addressed^3^. Yet, even if therapeutics against the existing epidemic strains are identified, there are several hundred other coronaviruses in active circulation within animal populations, many with the theoretical potential to infect humans. To help identify therapeutics for the current epidemic along with preparing for the next, there is a need for readily deployable small molecule screening assays that enable the identification of therapeutics that are broad-acting across a large collection of coronavirus strains.

During coronavirus infection, the RNA genome is delivered into cells and translated into a pair of polyproteins^4^. These polyproteins are then processed by a set of virally encoded proteases, of which the three-chymotrypsin-like protease (3CLpro) performs the majority of cleavage events^4^. As a result of its essential role in viral replication and high degree of conservation across all coronaviruses, 3CLpro enzymes represent important targets for therapeutic drug development^5,6^. Previous work expressing a variety of viral proteases within yeast and mammalian cellular systems have shown that protease expression can lead to profound cellular toxicity, which can be rescued by the addition of protease inhibitors^7–12^. We hypothesized that if the expression of coronavirus 3CLpro enzymes within mammalian cells leads to a similar toxic phenotype, this could form the basis of an easily implemented mammalian cell-based assay to evaluate protease inhibitors. While multiple assays exist to evaluate protease inhibitors, an assay of the nature we envisioned has clear advantages, as it requires minimal upfront cost or effort, is accessible to many biomedical research labs, does not involve the use of live virus, and requires no specialized reporter to read out protease activity. In contrast, *in vitro* protease assays using purified protein have formed the backbone of inhibitor screening, but require upfront efforts to isolate the pure protease and are not conducted under physiologic cellular conditions^13,14^. In addition, if one desires to identify broad-acting coronavirus inhibitors, one must purify and identify experimental conditions suitable for testing each protease *in vitro*. An alternative approach for identifying protease inhibitors is the use of live virus which is performed under more biologically relevant conditions, assuming relevant host cell systems can be identified, but requires intensive safety training and specialized biosafety protocols^15^. For many coronaviruses, no live virus assay exists, limiting the ability to test compounds within mammalian cell systems to a small subset of all coronaviruses^16^. Furthermore, compounds with activity against live virus may function through a number of mechanisms other than protease inhibition which cannot be readily determined, and may lead to undesired off-target activities which are not realized until much later in the drug development process^17,18^.

Here, we report a mammalian cell-based assay to identify coronavirus 3CLpro inhibitors that does not require the use of live virus. We demonstrate the utility of the assay using the SARS-CoV-2 3CLpro, with EC_50_ values obtained from the assay showing good concordance with traditional live virus testing for multiple compounds. We next establish the generality of the approach by testing a diverse set of 3CLpro enzymes. Finally, we perform a small molecule screen, along with structure-activity profiling of a set of compounds to find those with enhanced antiviral activity. The presented data support the use of our assay system for the discovery of small molecule 3CLpro inhibitors and the rapid characterization of their activity across multiple coronaviruses to identify those with broad inhibitory activity.

## Results

### Expression of the SARS-CoV-2 3CLpro in HEK293T cells results in cell toxicity and can be used as a readout of drug activity

Prior work has shown that the exogenous expression of viral proteases within cellular systems can result in growth suppression or cell death. Motivated by this work, we sought to determine the effect of expressing the SARS-CoV-2 3CLpro in HEK293T cells. Utilizing a cost-effective crystal-violet-based approach to quantify cell abundance, we observed that expression of SARS-CoV-2 3CLpro results in significant growth inhibition as compared to the expression of a control protein, enhanced yellow fluorescent protein (EYFP) (Fig. 1a-b)^19^. This suppression of growth was dependent upon the catalytic function of the enzyme, as mutating cysteine 145, which is essential for the enzyme’s peptidase activity, abolished the growth defect (Fig. 1a-b). We sought to determine if the observed growth defect could be rescued by incubating cells with GC376, a feline coronavirus inhibitor that was recently reported to have activity against SARS-CoV-2 3CLpro^20^. In comparison to untreated control cells, the addition of GC376 led to a robust increase in cell growth (Fig. 1c-d).

**Fig. 1.**
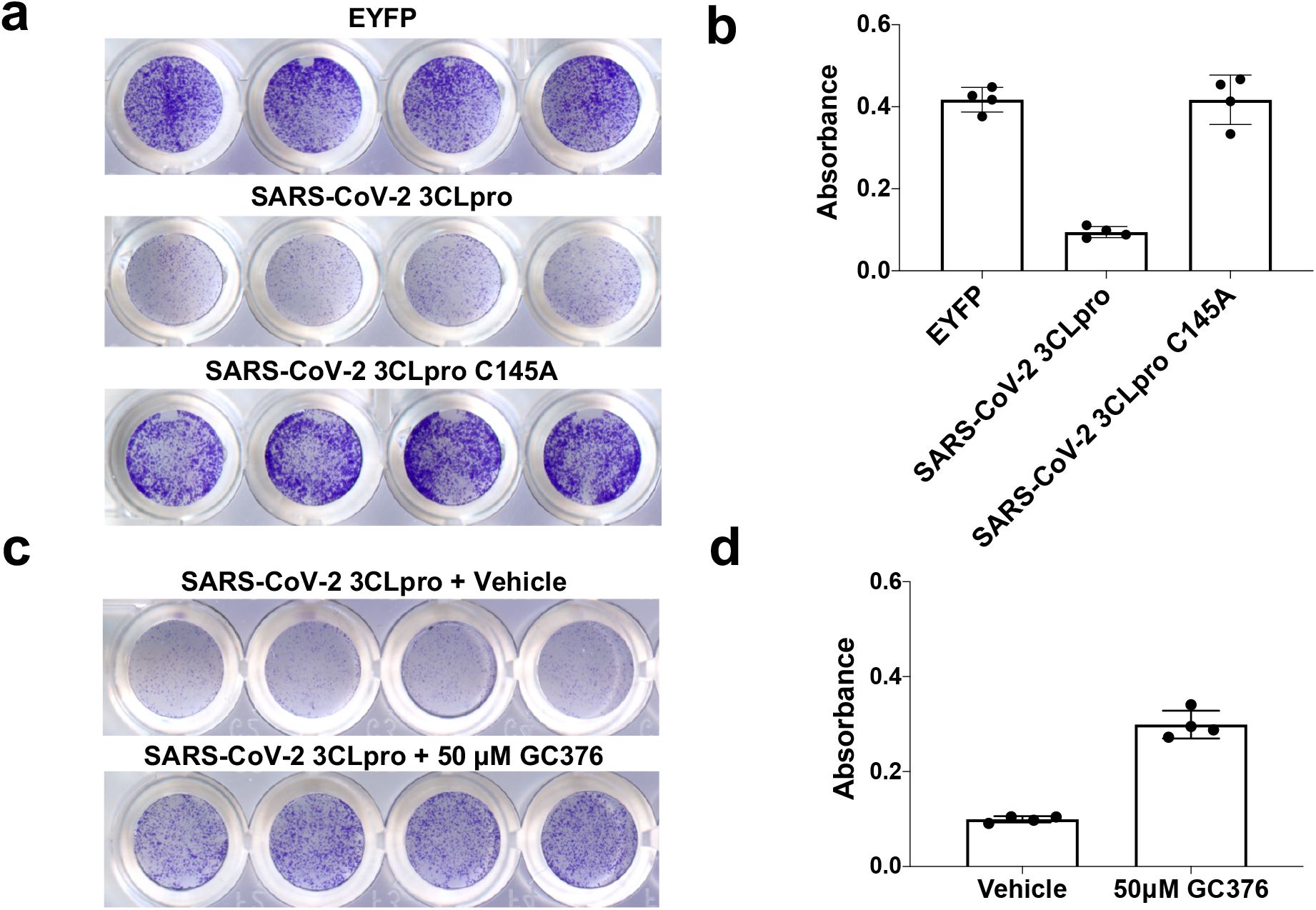
Expression of SARS-CoV-2 3CLpro in HEK293T cells results in toxicity that can be rescued by GC376. **a**. SARS-CoV-2 3CL toxicity is dependent on protease activity and can be visualized with crystal violet staining. **b**. Quantification of crystal violet staining in a. **c**. Treatment of SARS-CoV-2 3CLpro expressing cells with protease inhibitor GC376 results in rescue of cytotoxicity. **d**. Quantification of c. Data are shown as mean ± s.d. for four technical replicates.

### Compound rescue of transfected 3CLpro cytotoxicity is similar to results obtained with live virus

We next tested if this transfection-based assay could be used to determine compound EC_50_ values and whether the values showed any correlation with those obtained with live virus. After incubating SARS-CoV-2 3CLpro transfected cells with a range of GC376 concentrations, we calculated an EC_50_ of 3.30 μM, which is similar to published values reported using live virus on Vero E6 (EC_50_ 2.2 μM, 0.9 μM, 0.18 μM, 4.48 μM) and Vero 76 cells (EC_50_ 3.37 μM) (Fig. 2a and Table 1)^20–24^. We investigated the assay’s tolerance to deviation by varying the amount of plasmid transfected or the number of cells seeded into wells containing compound (Supplementary Fig. 1). In all cases, the assay was robust to variation, delivering a similar EC_50_ for GC376 across all conditions. We also tested an orthogonal method of quantifying cell abundance based on fluorescence microscopy and observed agreement with the results obtained with crystal violet staining (Supplementary Fig. 2). This suggests that the assay system is robust to variation and provides consistent results.

**Table 1.** Comparison of literature reported live virus based EC_50_ values compared to values generated during this study. CPE = Cytopathic effect.

| Protease | Drug | Calculated<br>EC<sub>50</sub> (μM) | Literature<br>Reported<br>Value (μM) | Method | Cell<br>Line | Citation |
| --- | --- | --- | --- | --- | --- | --- |
| SARS-CoV-2<br>3CLpro | GC376 | 3.3 | 3.37 | CPE | Vero 76 | Ma et. al. |
|  |  |  | 0.9 | Plaque<br>Assay | Vero E6 | Vuong et. al. |
|  |  |  | 0.18 | qPCR | Vero E6 | Luan et. al. |
|  |  |  | 2.2 | qPCR | Vero E6 | Froggatt et. al. |
|  |  |  | 4.48 | CPE | Vero E6 | Iketani et. al. |
| SARS-CoV-2<br>3CLpro | 11a | 6.89 | 2.06 | CPE | Vero E6 | This study |
|  |  |  | 0.53 | Plaque<br>Assay | Vero E6 | Dai et. al. |
| SARS-CoV-2<br>3CLpro | Compound<br>4 | 0.98 | 3.023 | CPE | Vero E6 | Iketani et. al. |
| SARS-CoV-2<br>3CLpro | GRL-0496 | 5.05 | 9.12 | CPE | Vero E6 | This study |
| SARS-CoV<br>3CLpro | GRL-0496 | 7.84 | 6.9 | CPE | Vero E6 | Ghosh et. al. |
| SARS-CoV-2<br>3CLpro | GC373 | 2.8 | 1.5 | Plaque<br>Assay | Vero E6 | Vuong et. al. |
|  |  |  | 4.83 | CPE | Vero E6 | This Study |

**Fig. 2.**
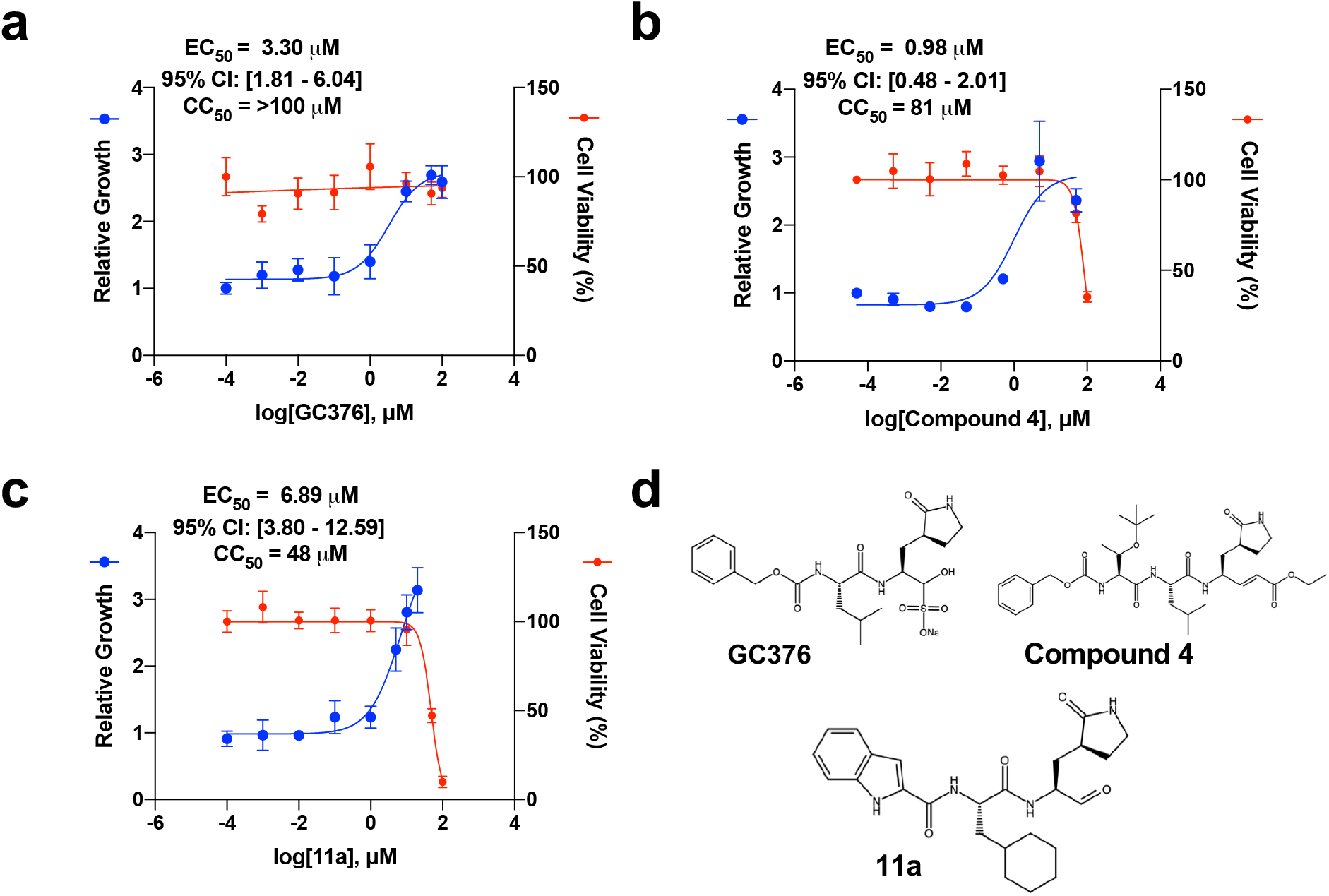
Dose response curves for SARS-CoV-2 3CLpro can be conducted with transfection-based assays. **a-c**. SARS-CoV-2 3CLpro can be inhibited by known 3CLpro inhibitors GC376, compound 4, and 11a. The toxicity of each compound was determined by treating EYFP-transfected cells with indicated concentrations of drug and is reported as Cell Viability. **d.**Chemical structures for each of the compounds tested. EC_50_ values are displayed as best-fit value alongside 95% confidence interval. CC_50_ values are displayed as best-fit value. Data are shown as mean ± s.d. for four technical replicates.

We next conducted dose-response profiling for two additional SARS-CoV-2 3CLpro inhibitors, compound 4 and compound 11a, and observed reversal of the toxic effect of the protease in a dose-dependent manner (Fig. 2b-c)^25,26^. In agreement with the results obtained with GC376, the EC_50_ value for compound 4 was comparable to those obtained with live virus, 0.98 μM and 3.023 μM, respectively (Table 1)^24^. Unexpectedly, we calculated an EC_50_ of 6.89 μM for 11a, which is approximately 10-fold higher than the literature reported value of 0.53 μM, based on viral plaque assay^26^. We have noticed that literature reported EC_50_ values from live virus testing could range over an order of magnitude depending on the exact method employed, as is the case for GC376 (Table 1). To resolve this discrepancy between the transfection-based approach and the live virus assay, we conducted live virus testing of 11a using the commonly employed readout of cytopathic effect in Vero E6 cells and observed closer concordance with our transfection-based results (Supplementary Figure 3 and Table 1)^20,24,27^. To measure the toxicity of each compound, we exposed EYFP-transfected cells to each molecule and determined CC_50_ values (Fig. 2). We also calculated the selectivity index (SI) for each compound tested in this study (Supplementary Table 1).

We hypothesized that the assay would be able to distinguish between compounds that are only active on the purified SARS-CoV-2 3CLpro and those that are able to inhibit the live virus through protease inhibition. In general, we observe concordance between compounds showing activity within this transfection-based 3CL assay and live virus studies (Supplementary Fig. 4a-e)^13^. However, within the assay, we did not observe activity for ebselen, a small molecule with demonstrated activity against the SARS-CoV-2 3CLpro *in vitro* and activity against the SARS-CoV-2 live virus (Supplementary Fig. 4f). We suggest that this may be due to ebselen targeting more than 3CLpro within the live virus assay, which is in line with work showing that ebselen is highly reactive and readily forms selenosulfide bonds with numerous proteins including the SARS-CoV-2 papain-like protease (PLP)^18,28,29^.

**Fig. 4.**
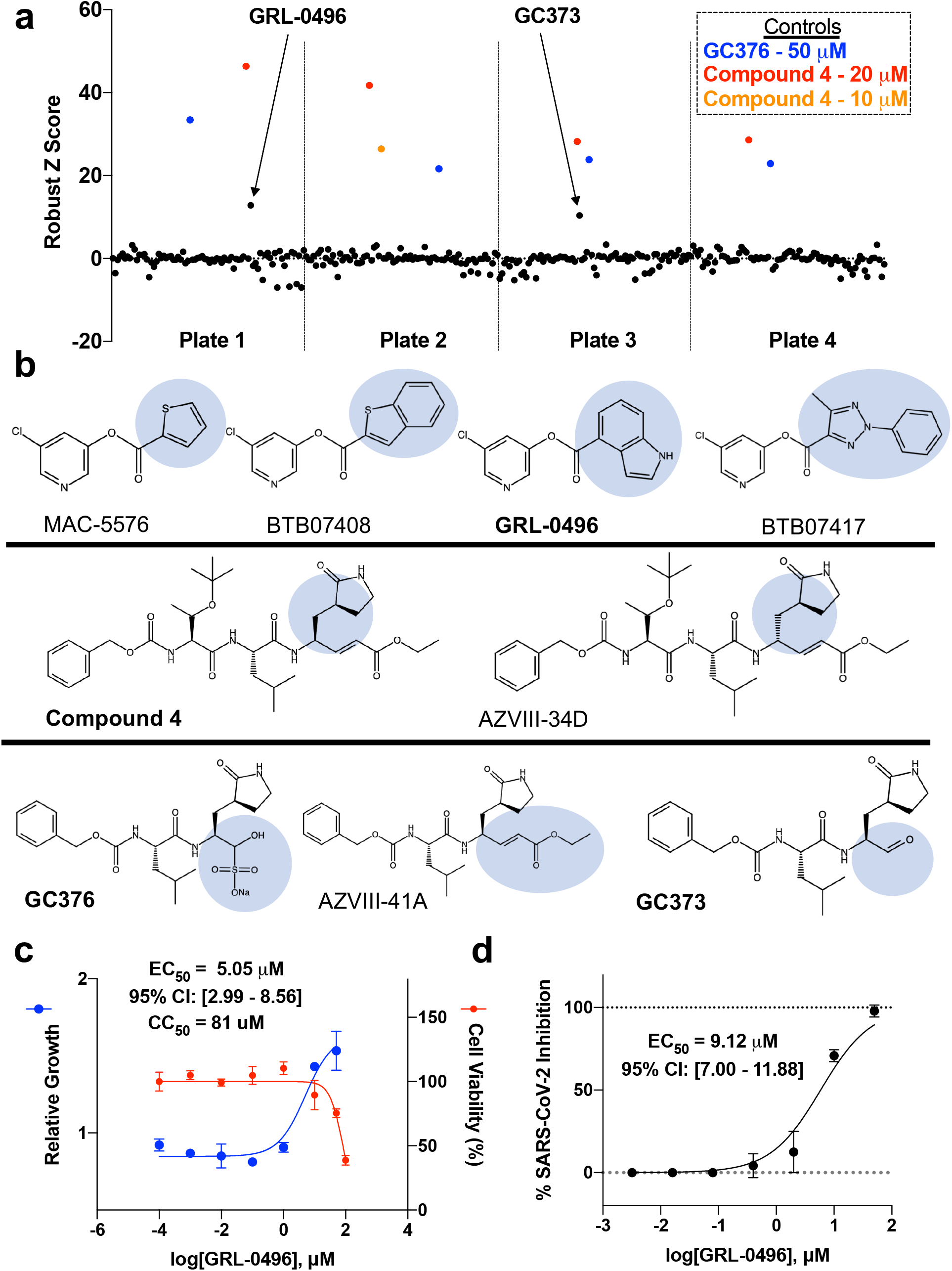
Small-scale drug screen and structure-activity profiling at 10 μM identify two compounds, GC373 and GRL-0496, with activity against the SARS-CoV-2 3CLpro. **a**. Identification of hits from the drug screen and structure-activity profiling. Positive control compounds were included in each plate and are highlighted. **b**. Compounds with structural similarity to known inhibitors. Compounds in bold are molecules that show activity against the SARS-CoV-2 3CLpro at 10 μM. **c**. Dose-response profiling and cytotoxicity determination of GRL-0496 against the SARS-CoV-2 3CLpro. **d**. Live virus testing of GRL-0496 against SARS-CoV-2. EC_50_ values are displayed as best-fit value alongside 95% confidence interval. The live virus assay was conducted with two biological replicates, each with three technical replicates and the EC_50_ value was derived from all replicates. CC_50_ values are displayed as best-fit value. Data are shown as mean ± s.d. for three or four technical replicates.

### Demonstrating assay compatibility across a range of coronavirus 3CLpro enzymes

We hypothesized that this assay may be used to study other coronavirus 3CLpros to enable users to identify broad-acting inhibitors, as constructs containing other 3CLpro enzymes could be readily synthesized. To test the assay’s generality, we created expression constructs for 3CL proteases from five other coronaviruses (SARS-CoV, MERS-CoV, Bat-CoV-HKU9, HCoV-NL63 and IBV) with variable amino acid identity compared with SARS-CoV-2 3CLpro (Supplementary Fig. 5a). For each of these proteases, we confirmed that expression in mammalian cells resulted in toxicity that is dependent upon the enzyme’s catalytic activity (Supplementary Fig. 5b). Next, we tested GC376, compound 4, and 11a across this panel of proteases. GC376, a drug originally identified for use against the Feline Infectious Peritonitis virus, showed EC_50_ <10 μM for the most, but not all of proteases tested^30^. Unexpectedly, compound 4, which was originally designed as a SARS-CoV 3CLpro inhibitor showed particular potency against IBV 3CLpro (EC_50_ = 0.058 μM) along with broad activity (EC_50_ <10 μM) for all other 3CL proteases tested. In contrast to GC376 and compound 4, 11a had a relatively narrow activity spectrum with EC_50_ <10 μM against only SARS-CoV and SARS-CoV-2 3CLpro enzymes (Fig. 3). Of note, in all cases where previous live virus data was available, the EC_50_ values obtained from this transfection-based assay were similar (Table 1).

**Fig. 3.**
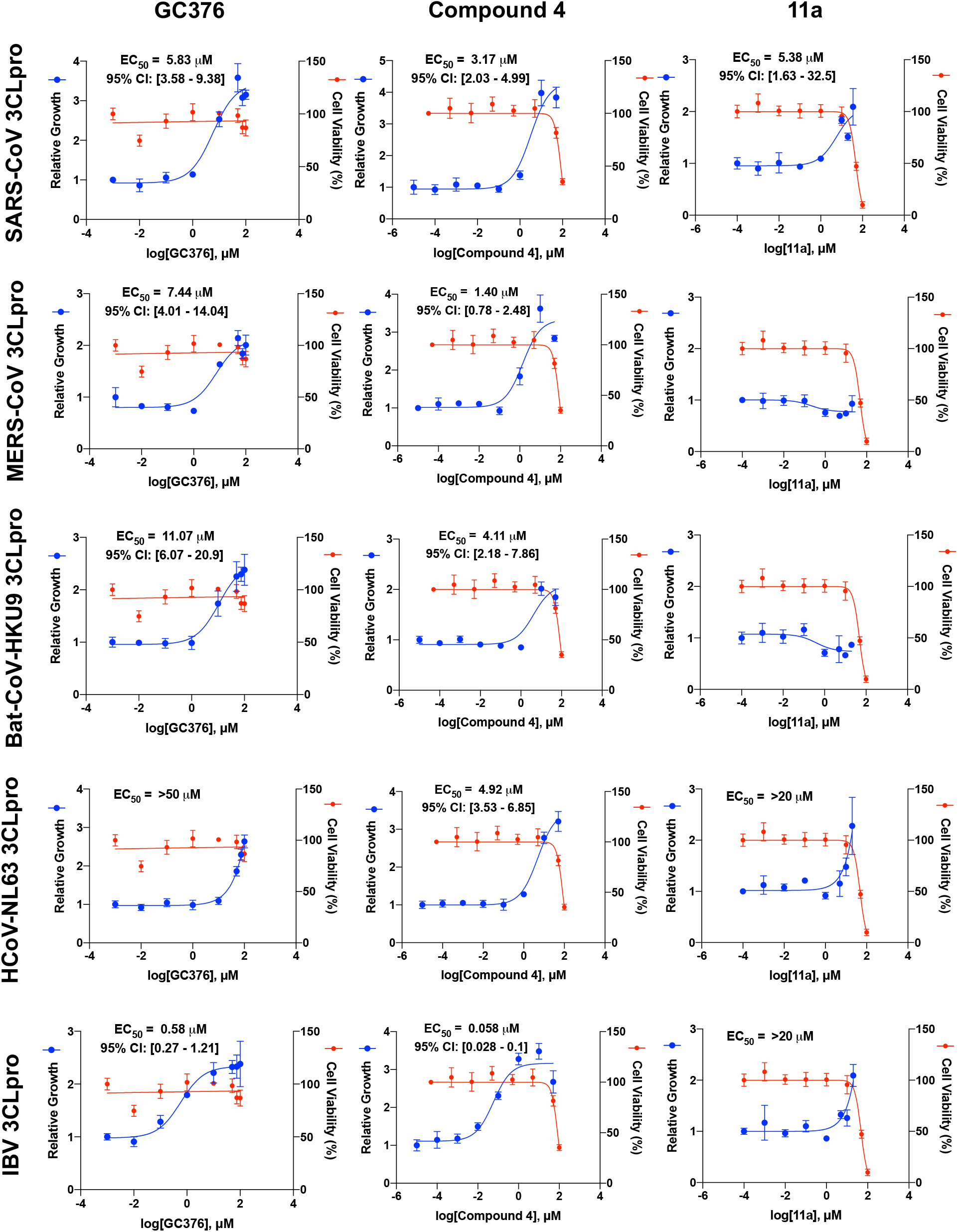
The activity of GC376, compound 4, and 11a show variable effectiveness and potency against the coronavirus 3CL proteases from SARS-CoV, MERS-CoV, Bat-CoV-HKU9, HCoV-NL63, and IBV. EC_50_ values are displayed as best-fit value alongside 95% confidence interval. CC_50_ values are displayed as best-fit value. Data are shown as mean ± s.d. for three or four technical replicates.

### Rapid testing to identify SARS-CoV-2-3CL protease inhibitors

Having further determined the assay’s ability to examine the effects of active individual compounds, we sought to determine its suitability for small molecule screening. Before performing the screen, we optimized the testing parameters to ensure suitable performance characteristics (Supplementary Fig. 6 and Methods)^31^. We compiled a collection of 162 diverse protease inhibitors, along with compounds with reported *in vitro* activity against 3CLpro enzymes or structural similarity to known 3CLpro inhibitors (Supplementary Table 2 and Supplementary Table 3). Of the nearly 200 compounds tested, two potent hits were identified, GC373 and GRL-0496 (Fig. 4a, Supplementary Table 2)^32^.

We noted that GC373 is structurally similar to its prodrug GC376, except for the change of the bisulfide salt adduct to an aldehyde warhead^22,33^. Additional testing of GC373 revealed it to have a similar EC_50_ as GC376 in both the transfection assay and when tested against live SARS-CoV-2 virus, suggesting that the differences in structure has a minimal effect on their potency (Supplementary Fig. 7 and Table 1), although solubility may be affected^33^. The other hit from the screen, GRL-0496, shares structural similarity to several other compounds within the library, one of which is a previously reported 3CLpro inhibitor (MAC-5576) that failed to show activity against SARS-CoV-2 in a live virus assay^24,34^. Additional testing of GRL-0496 revealed it to have an EC_50_ of 5.05 μM against SARS-CoV-2 3CLpro within our transfection-based assay (Fig. 4c). To verify GRL-0496’s activity, we tested it against live SARS-CoV-2 virus, and confirmed its potency (EC_50_ = 9.12 μM) (Fig. 4d). We next tested GRL-0496 against the full panel of 3CLpro enzymes we had previously examined and observed a narrow range of activity, with EC_50_ <10 μM only observed against SARS-CoV 3CLpro and SARS-CoV-2 3CLpro, in agreement with previous live virus testing (Supplementary Fig. 8 and Table 1)^27^. We also tested GC373 against the full panel of 3CLpro enzymes and observed concordance with GC376, with SARS-CoV 3CLpro, MERS-CoV 3CL pro, and IBV 3CL pro demonstrating an EC_50_ <10 μM (Supplementary Fig. 8).

## Discussion

Given the potential for protease inhibitors in the treatment of viral illnesses, small molecule inhibitors of coronavirus 3CL proteases represent a promising avenue for treating infections caused by this large family of viruses. Here, we present a simplified assay to identify candidate inhibitors under physiologic cellular conditions. This approach presents significant advantages over other methods to detect 3CL protease inhibitory activity with its ease of use and ability to be performed with equipment and reagents commonly available to many biomedical research laboratories. While conventional methods for identifying 3CL protease inhibitors make use of *in vitro* purified protease, the isolation of sufficiently pure enzyme in its native state can be costly and labor intensive. Furthermore, assays using purified protease fail to consider cell permeability and the influence of the extracellular and intracellular milieu on compound activity. In comparison to live virus-based assays, the outlined approach does not require extensive biosafety containment. These data also suggest that the approach described here is applicable to a number of coronaviruses for which live virus assays may not be available or would be deemed ethically challenging to be performed even with extensive biosafety infrastructure^35,36^. Finally, because the phenotype assayed within this approach is driven solely by protease activity, it may enable the distinction between compounds with multiple biological targets and subsequent potential for off-target toxicity from those that function primarily as 3CLpro inhibitors.

A number of surrogate assays that aim to identify molecules with activity against proteins encoded by SARS-CoV-2 have recently been reported^37–39^. These assays were developed to provide physiologically meaningful molecule testing without needing to use live SARS-CoV-2 cultures in order to allow for rapid testing and widespread adoption to labs without necessary safety infrastructure. The aforementioned assays have focused primarily on identifying neutralizing antibodies targeted at the SARS-CoV-2 spike (S) protein. To our knowledge, a reporter-free surrogate assay to identify coronavirus 3CL protease inhibitors validated with multiple known coronavirus 3CL protease inhibitors has not been reported.

Within the literature, EC_50_ values obtained for a 3CLpro inhibitor against live virus can show a broad range of potency, with some compounds demonstrating EC_50_ values across multiple orders of magnitude (Table 1). These differences appear to be driven by variation in experimental setup such as cell line used, assay readout, incubation period, and initial concentration of virus added. While we have observed agreement between the EC_50_ values obtained from the described transfection-based method and those reported in the literature, given the differences in EC_50_ across assays, we suggest caution when comparing results across studies. By developing this transfection-based 3CLpro testing platform, we hope to facilitate the discovery of new coronavirus inhibitors while also facilitating the comparison of existing inhibitors within a single simplified assay system. Furthermore, we propose that this cellular protease assay system could be industrialized to screen and optimize a large number of compounds to discover potential treatments for future viral pandemics.

## Methods

### Cell Lines and Cell Culture

HEK293T and HEK293 cells used in this study were obtained from ATCC. Cells were maintained at 37°C in a humidified atmosphere with 5% CO_2_. HEK293T and HEK293 cells were grown in Dulbecco’s Modified Eagle Medium (DMEM, Invitrogen) which was supplemented with 10% fetal bovine serum (Gibco) and penicillin-streptomycin (Invitrogen). HEK293T and HEK293 cells were confirmed to be free of mycoplasma contamination with the Agilent MycoSensor PCR Assay Kit. To obtain HEK293 cells stably expressing EYFP, cells were co-transfected with EYFP plasmids harboring the *piggyBac* transposon (pPB bsr2-EYFP) (Yusa et al., 2009) and pCMV-mPBase (mammalian codon-optimized PBase) encoding a piggyBac transposase using Lipofectamine 2000 (Invitrogen) according to the manufacturer’s instructions. One day after transfection, the transfected cells were selected with 10 μg/mL of blasticidin (Invitrogen).

### Transfections and Drug Selections

24 h prior to transfection, 293T cells were seeded at 65-75% confluency into 24-well plates coated for 30 min with a 1 mg/mL solution of poly-D-lysine (MP Biomedicals Inc.) and washed with PBS (Gibco) once prior to media addition. The next day, 500 ng of 3CLpro expression plasmid, unless otherwise stated, was incubated with Opti-MEM and Lipofectamine 2000 for 30 min at room temperature prior to dribbling on cells, as per manufacturer’s protocol. For plating into drug conditions, 20 h after transfection, cells were washed once with PBS and 200 μL Trypsin-EDTA 0.25% (Gibco) was added to cells to release them from the plate. Trypsinized cell slurry was pipetted up and down repeatedly to ensure a single cell suspension. 96-well plates were coated with poly-D-lysine, either coated manually with 1 μg/mL poly-D-lysine in PBS solution for 30 min or purchased pre-coated with poly-D-lysine (Corning). Wells were filled with 100 μL of media ± drug and 1 μg/mL puromycin and were seeded with 9 μL of trypsinized cell slurry. For higher throughput experiments, multiple individually transfected wells of a 24-well plate were combined after trypsinization and prior to seeding in drug. After seeding into wells containing drug and puromycin, cells were incubated for 48 h unless otherwise specified.

### Plasmids

All vectors used in this study were cloned into the pLEX307 backbone (Addgene #41392) using Gateway LR II Clonase Enzyme mix (Invitrogen). 3CL proteases used in this study were generated using gene fragments ordered from Twist Biosciences. Inactive 3CL proteases were generated by site directed mutagenesis of the essential catalytic cysteine. DNA was transformed into NEB 10-beta high efficiency competent cells. Sanger sequencing to verify proper inserts were done for all plasmids used in this study (Genewiz).

Plasmid DNA was isolated using standard miniprep buffers (Omega Biotek) and silica membrane columns (Biobasic). To reduce batch-to-batch variability between plasmid DNA isolations and its subsequent impact on transfection efficiency, multiple plasmid DNA extractions were conducted in parallel, diluted to 50 ng/μL and pooled together.

### Crystal Violet Staining and Quantification

The crystal violet staining protocol was adapted from Feoktistova et. al.^19^ Briefly, after compound incubation with 3CLpro expressing cells in 96-well plates, the medium was discarded and cells were washed once with PBS. Cells were incubated with 50 μL of crystal violet staining solution (0.5% crystal violet in 80% water and 20% methanol) and rocked gently for 30 min. The staining solution was removed and cells were washed four times with water using a multichannel pipette. Stained cells were left to dry for ≥4 h on the laboratory bench or within the chemical hood. The crystal violet staining solution was eluted by the addition of 200 μL of methanol over the course of 30 min with gentle rocking. Plates were sealed with parafilm to mitigate methanol evaporation. 100 μL of eluted stain from each well was transferred to a new 96-well plate for reading in a Tecan Infinite F50 plate reader. Absorbance was measured at 595 nm twice and values were averaged between replicate measurements. Blank wells were included in each batch of experiments, and absorbance values were normalized by background levels of staining from blank wells.

### Statistical Analysis of Dose Response Curves

For analysis of crystal violet staining experiments, relative growth was calculated from background normalized absorbance values. Test wells containing drug were divided by average background normalized values from wells where cells were expressing protease and exposed to vehicle, when available. Otherwise, values were normalized by values from protease-expressing cells exposed to the lowest concentration of drug included in the dose-response curve. When there were significant deviations from protease-expressing cells exposed to no drug and protease-expressing cells exposed to lowest concentrations of drug included in the dose-response curve, experiments were repeated with normalization by protease-expressing cells exposed to no drug. CC_50_ values were calculated in Prism using the nonlinear regression functionality and derived from dose-response curves with EYFP transfected cells. A nonlinear curve fitting function accounting for variable curve slopes was calculated by plotting the normalized response as a function of log(compound). Similarly, EC_50_ values were calculated in GraphPad Prism also using the nonlinear regression functionality. A nonlinear curve fitting function measuring the stimulatory response of a compound as a function of an unnormalized response was used to calculate the EC_50_. All reported values were confirmed to not have ambiguous curve fitting. The 95% confidence interval of EC_50_ calculations was also calculated and included.

For analysis of live virus experiments, EC_50_ values were determined by fitting a nonlinear curve to the data with the assumption of a normalized response (GraphPad Prism). Cells were confirmed as mycoplasma negative prior to use.

### Compound Screening

For screening condition optimization, we measured the Z-Factor for replicates of positive controls GC376, tested at 50 μM, and compound 4, tested at 20 μM. Replicate measurements were recorded for DMSO negative controls and positive control compounds after 48, 72, and 96 h of incubation with drug. Background normalized crystal violet absorbance values at each timepoint were collected.

During the drug screen, within each of the four plates screened, two positive controls wells were included to ensure assay reliability, along with several wells with the negative control 0.1% DMSO condition. All compounds were screened at 10 μM resuspended in DMSO (Fisher Scientific). For hit selection, we employed a robust z-score method. We first normalized data using a robust z-score that uses median and median absolute deviation (MAD) instead of mean and standard deviation. We then used a threshold of 3.5 MAD to determine which drugs rescued the cytotoxicity imposed by expression of the viral protease^40^.

### Live Virus Assay

The SARS-CoV-2 strain 2019-nCoV/USA_WA1/2020 was grown and titered in Vero-E6 cells. One day before the experiment, Vero-E6 cells were seeded at 30,000 cells/well in 96 well-plates. Serial dilutions of the test compound were prepared in media (EMEM + 10% FCS + penicillin/streptomycin), pipetted onto cells, and virus was subsequently added to each well at an MOI of 0.2. Cells were incubated at in a humidified environment at 37 °C with 5% CO_2_ for 72 h after addition of virus. Cytopathic effect was scored by independent, blinded researchers. The reported cytopathic effect value represents the average from two independent reviewers. Percent Inhibition was calculated by comparison to control wells with no inhibitor added. All live virus experiments were conducted in a biosafety level 3 lab.

### Microscopy

Cells plated on a 96-well plate (Greiner Bio-OneTM) and washed once immediately before imaging. EYFP fluorescence imaging was performed using an Axio Observer 7 microscope (Zeiss) equipped with a Plan-Apochromat 10X objective (0.45 N.A.) with 1-by-1 pixel binning. Optical Illumination bias was empirically derived by sampling background areas and subsequently used to flatten images. After a global background subtraction, cell density was calculated based on area of EYFP intensity.

### Compounds and Chemical Synthesis

GC376 was purchased from Aobious. Myrecetin, rupintrivir, grazoprevir, saquinavir, fosamprenavir, indinavir, apigenin, quercetin, famotidine, MDL28170, bicailein, betrixaban, and amentoflavone were purchased from Fisher Scientific. Tipranavir was purchased from Cayman Chemical. MAC5576, MAC22272, MAC8120, MAC30731, BTB07404, BTB07408, MWP00332, BTB07417, MWP00508, MWP00333, BTB07407, SPB08384, SPB06613, SPB06636, SPB06591, SPB06593, MWP00709, CC42746, BTB07789, BTB07420, MWP00710, BTB07421, SCR00533, and SEW03089 were purchased from Maybridge. GRL-0496 and GRL-0617 were purchased from Focus Biomolecules. AZVIII-40A (1,2-Benzisothiazol-3(2*H*)-one) was purchased from Alfa Aesar. Other protease inhibitors in table below were purchased from TargetMol.

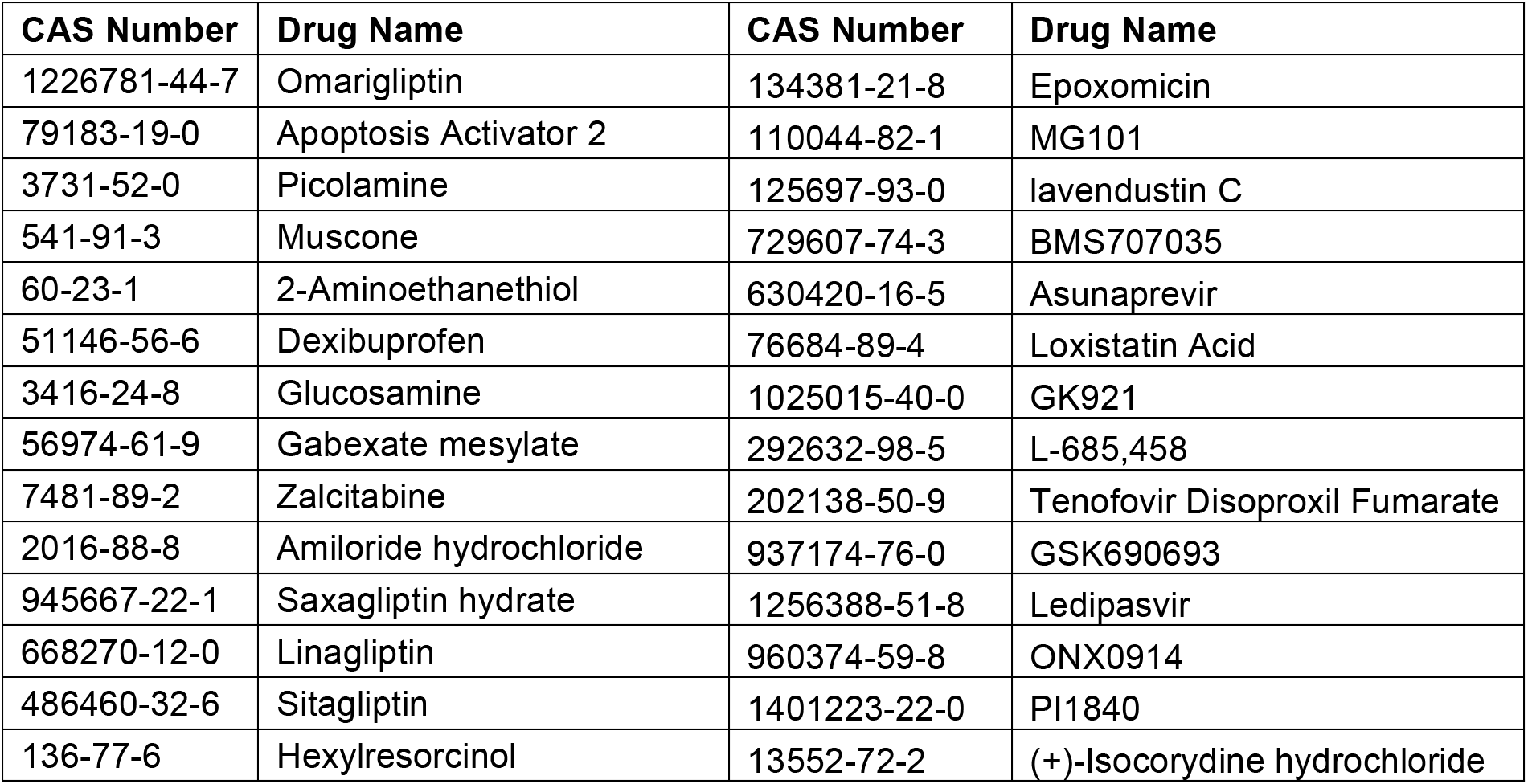
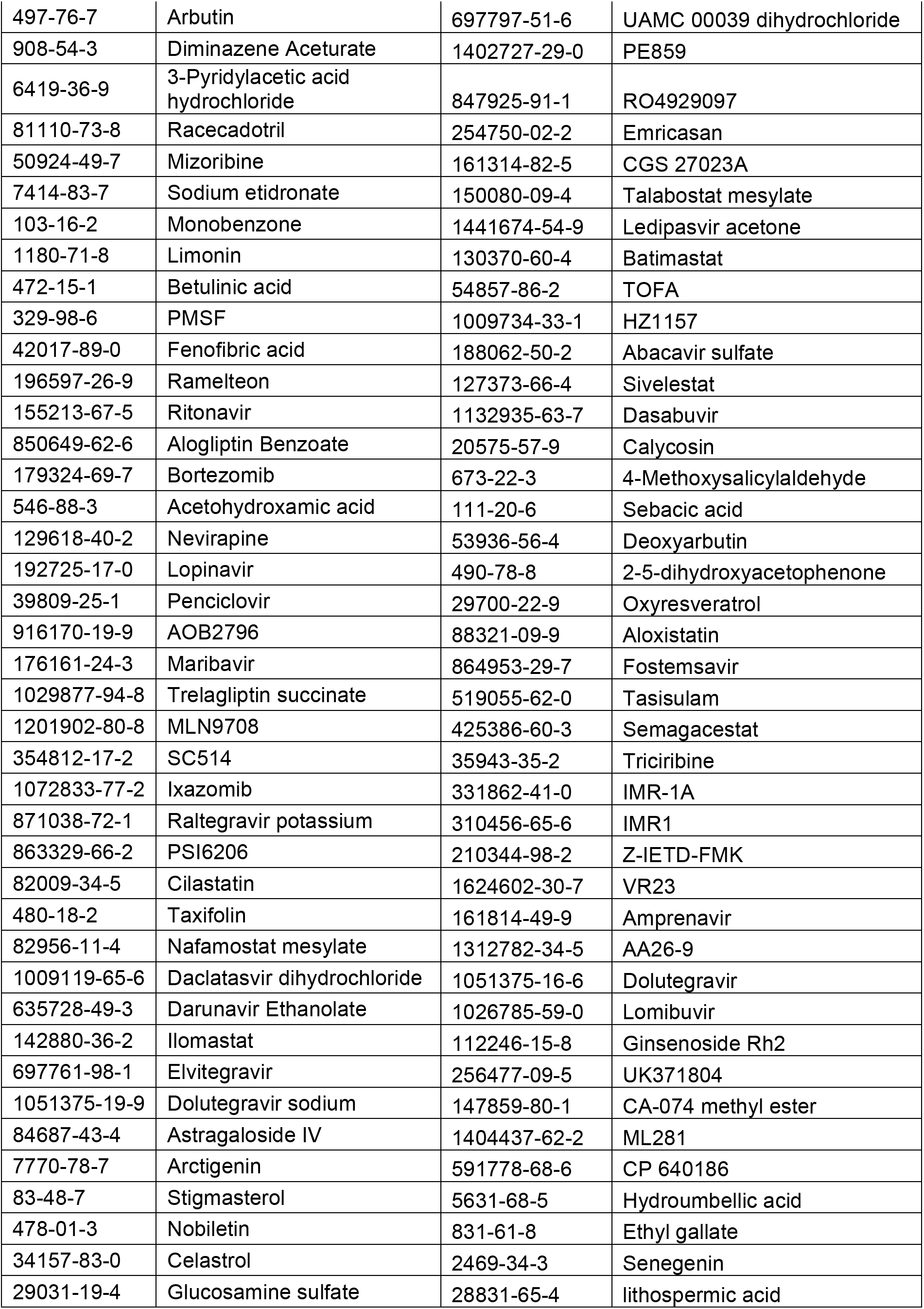
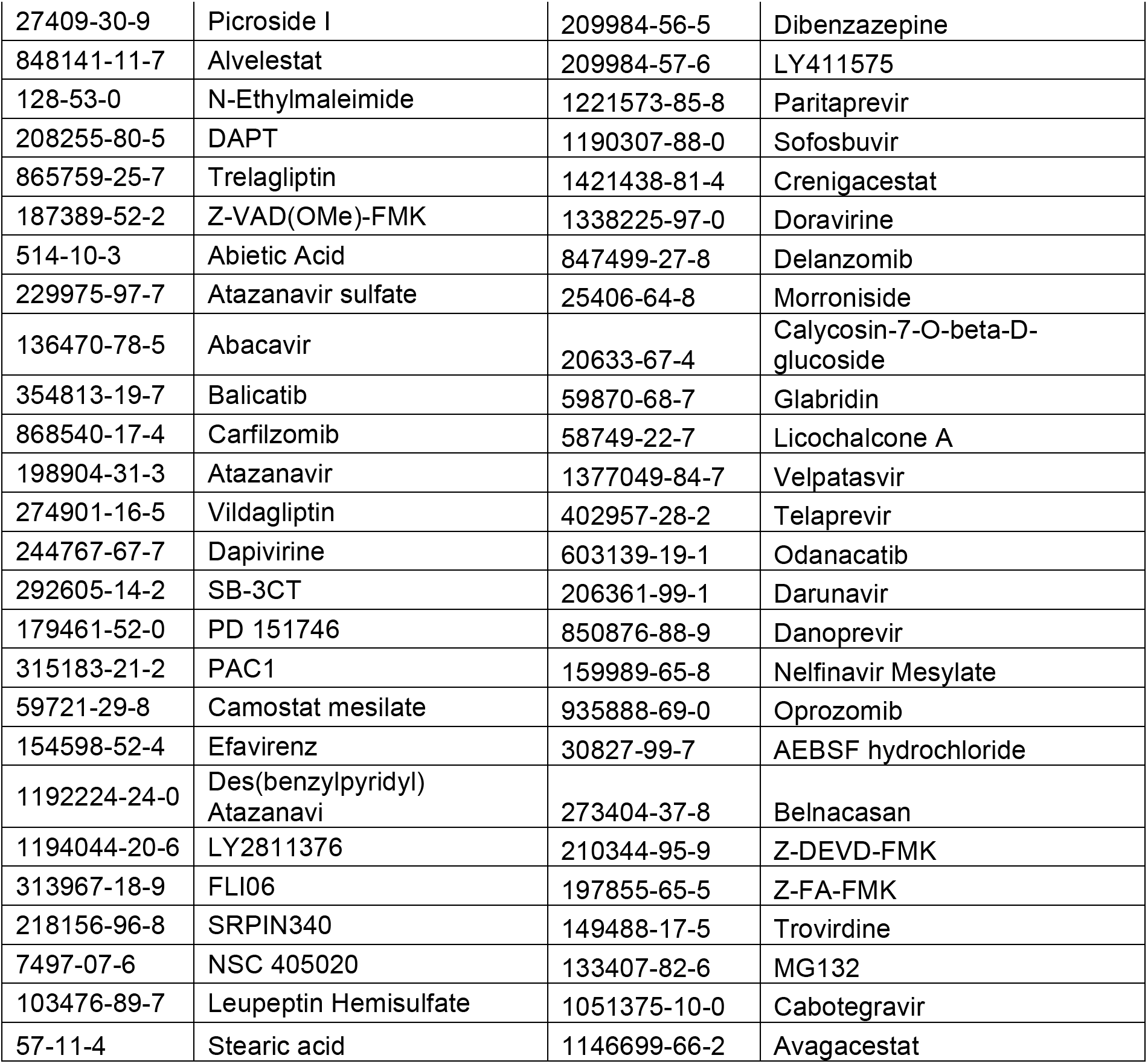

All other compounds used in the study were synthesized and quality checked according to the following protocols.

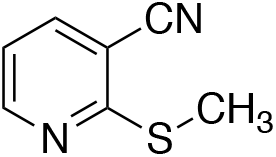

2-(methylthio)nicotinonitrile (xx-1) The title compound was prepared according to a published procedure; spectral data are in agreement with literature values^41^.

^1^H NMR (500 MHz, CDCl_3_) δ 8.60 (dd, *J* = 5.0, 1.8 Hz, 1H), 7.79 (dd, *J* = 7.7, 1.8 Hz, 1H), 7.07 (dd, *J* = 7.7, 4.9 Hz, 1H), 2.64 (s, 3H).

^13^C NMR (126 MHz, CDCl_3_) δ 163.7, 152.2, 140.6, 118.4, 115.7, 107.5, 13.4.

HRMS High accuracy (ASAP): Calculated for C_7_H_6_N_2_S (M+H)^+^: 151.0330; found: 151.0327.

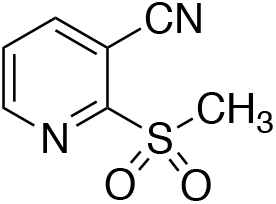

2-(methylsulfonyl)nicotinonitrile (xx-2) The title compound was prepared from xx-1 according to a modified procedure from the literature^42^.

Pyridine xx-1 (87 mg, 0.58 mmol, 1 equiv) was weighed into 50 mL round bottom flask equipped with a stir bar. The solid was dissolved in 10 mL of anhydrous MeOH, followed by portion-wise (usually 3 portions) additions of mCPBA (500 mg, 2.9 mmol, 5 equiv). Substrate conversion was monitored via TLC analysis (70% EtOAc:Hex to 100% EtOAc), with up to an additional 5 equiv of mCPBA added if necessary. Upon complete conversion of starting material, the reaction was quenched with 20 mL of saturated aqueous NaHCO_3_, diluted with an additional 20 mL of DCM. The layers were separated and the organic layer was further washed (2x) with 10 mL of of saturated aqueous NaHCO_3_. The organic layer was then dried *in vacuo* and purified via silica gel column chromatography (100% EtOAc to 5% MeOH:EtOAc) to yield 51 mg (48% yield) of the desired sulfone as a white solid.

^1^H NMR (400 MHz, CDCl_3_) δ 8.86 (dd, *J* = 4.8, 1.6 Hz, 1H), 8.26 (dd, *J* = 7.9, 1.6 Hz, 1H), 7.70 (dd, *J* = 7.9, 4.8 Hz, 1H), 3.39 (s, 3H).

^13^C NMR (101 MHz, CDCl_3_) δ 159.8, 151.8, 143.8, 126.8, 113.2, 107.4, 40.1.

HRMS High accuracy (ASAP): Calculated for C_7_H_6_N_2_O_2_S (M+H)^+^: 183.0228; found: 183.0223.

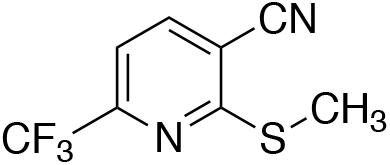

2-(methylthio)-6-(trifluoromethyl)nicotinonitrile (xx-3) The title compound was prepared following the published procedure, the product (yellow solid) was carried towards the synthesis of xx-4 without further purification^41^.

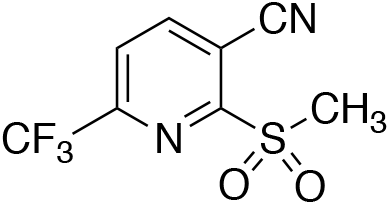

2-(methylsulfonyl)-6-(trifluoromethyl)nicotinonitrile (xx-4) The title compound was prepared from xx-3 according to a modified procedure from the literature^42^.

Pyridine xx-2 (105 mg, 0.50 mmol, 1 equiv) was weighed into 50 mL round bottom flask equipped with a stir bar. The solid was dissolved in 10 mL of anhydrous MeOH, followed by portion-wise (usually 3 portions) additions of mCPBA (0.86 mg, 5 mmol, 10 equiv). Substrate conversion was monitored via TLC analysis (40% EtOAc:Hex to 60% EtOAc). Upon complete conversion of starting material, the reaction was quenched with 20 mL of saturated aqueous NaHCO_3_, diluted with an additional 20 mL of DCM. The layers were separated and the organic layer was further washed (2x) with 10 mL of of saturated aqueous NaHCO_3_. The organic layer was then dried *in vacuo* and purified via silica gel column chromatography (40% to 80% EtOAc:Hex) to yield 38 mg (30% yield) of the desired sulfone as a white solid.

^1^H NMR (400 MHz, CDCl_3_) δ 8.49 (dd, *J* = 8.1, 0.7 Hz, 1H), 8.06 (d, *J* = 8.1 Hz, 1H).

^13^C NMR (101 MHz, CDCl_3_) δ 160.4, 149.7 (q, *J* = 37.6 Hz), 146.0, 123.7 (q, *J* = 2.3 Hz), 120.0 (q, *J* = 275.5 Hz), 112.1, 109.7, 39.7.

^19^F NMR (376 MHz, CDCl_3_) δ-68.28.

HRMS High accuracy (ASAP): Calculated for C_8_H_5_F_3_N_2_O_2_S (M+H)^+^: 251.0102; found: 251.0104.

Compound 11a was synthesized according to the specified protocol in Dai. et. al. 2020^26^. Compound 11a was confirmed by LCMS with m/z = 453 (M+1) and 451 (M-1).

Compound 4 was synthesized according to the procedure described in Yang et. al 2006^25^. AZVIII-34D was formed as a byproduct (15%) in the synthesis of Compound 4 and was isolated by RP HPLC. ^1^H NMR (400 MHz, CDCl_3_) 7.88 (s, 1H), 7.44-7.34 (m, 5H), 6.80 (d, 1H, J = 15.4 Hz), 6.69 (s, 1H), 6.16-5.87 (m, 2H), 5.12 (s, 2H), 4.76 (s, 1H), 4.42 (s, 1H), 4.31-3.99 (m, 4H), 3.39 (s, 2H), 2.40-1.45 (m, 8H), 1.29–1.20 (m, 12H), 1.13-1.02 (m, 3H), 1.00 – 0.89 (m, 6H). MS M+H = 631.

AZVIII-38, AZVIII-30, and AZVIII-42 were synthesized according the procedures described in Mou et. al. 2008^43^. AZVIII-37A was synthesized as described in Prior et al. 2013. AZVIII-33B was synthesized according to the method described in Amblard et. al. 2018^44^.

AZVIII-41A was synthesized according to the procedure described for synthesizing AZVIII-33B in Amblard et. al. 2018 and substituting Z-Leu-OH for BOC-Leu-OH^44^. ^1^H NMR (400 MHz, CDCl_3_) 7.77 (s, 1H), 7.41-7.27 (m, 5H), 6.80 (dd, 1H, J = 5.5, 15. 6 Hz), 5.98-5.84 (m, 2H), 5.15-5.03 (m, 2H), 4.55 (bd s, 1H), 4.27 (bd s, 1H), 4.23-4.07 (m, 3H), 3.38-3.23 (m, 2H), 2.56–1.40 (m, 8H), 1.33 – 1.20 (m, 3H), 0.95 (s, 6H). MS M+H = 474.

AZVII-43A was synthesized by treatment of a dichloromethane solution of AZVIII-30 and AZVIII-40A with triphenyl phosphine and diethyl azodicarboxylate. ^1^H NMR (400 MHz, CDCl_3_) 7.91 (d, 1H, J = 8.0), 7.76 (d, 1H, J = 8.0), 7.55-7.34 (m, 2H), 6.16 (bd s, 1H), 5.29 (bd s, 1H), 4.73-4.39 (m, 2H), 4.35-3.96 (m, 1H), 3.58-3.16 (m, 2H), 3.79-3.43 (m, 2H), 3.39-3.91 (m, 1H), 1.98-1.77 (m, 1H), 1.73-1.57 (m, 1H), 1.42 (s, 9H). MS M+H = 392.

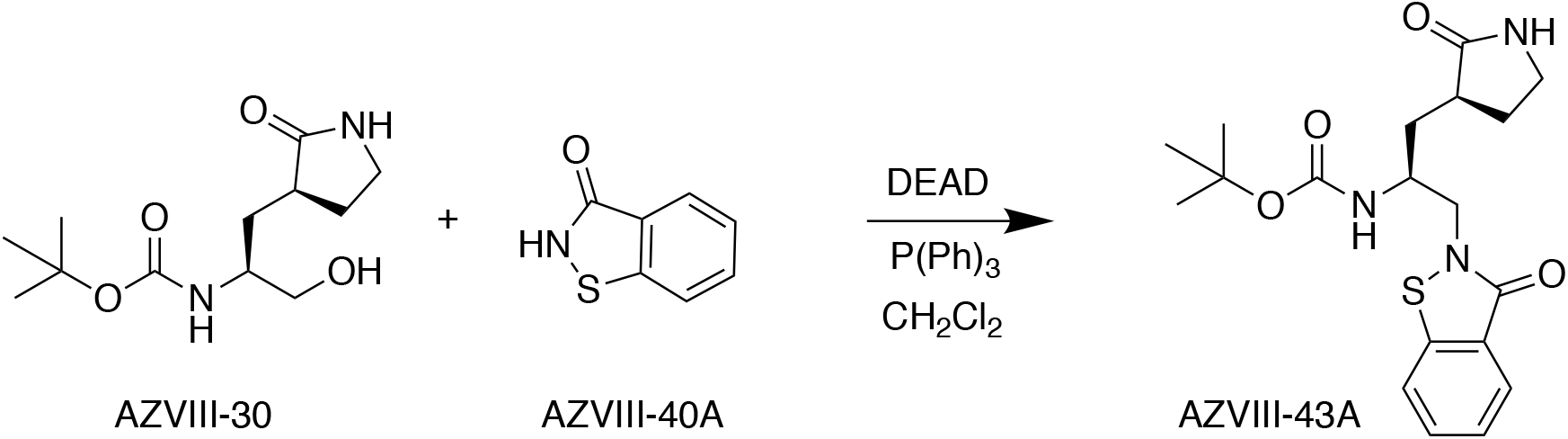

AZVIII-44B, AZVIII-44D, AZVIII-44E, AZVIII-44H, AZVIII-49C, and AZVIII-49F were synthesized by treatment of the appropriate arylalkylamine or heteroarylalkylamine with the corresponding arylsulfonyl chloride or heteroarylsulfonyl chloride and diisopropylethylamine in dichloromethane. AZVIII-57D was synthesized by treatment of AZVIII-44D with methyl bromoacetate and potassium carbonate in dimethylformamide to give the corresponding ester which was reduced to the corresponding alcohol by treatment with lithium borohydride in tetrahydrofuran. The alcohol was then treated with 1,1,1-tris(acetyloxy)-1,1-dihydro-1,2-benziodoxol-3-(1*H*)-one and sodium bicarbonate in dichloromethane to give to the corresponding aldehyde AZVIII-57D. AZVIII-57G was synthesized from *N*-benzyl-4-methoxybenzenesulfonamide according to the procedure used to synthesize AZVIII-57D.

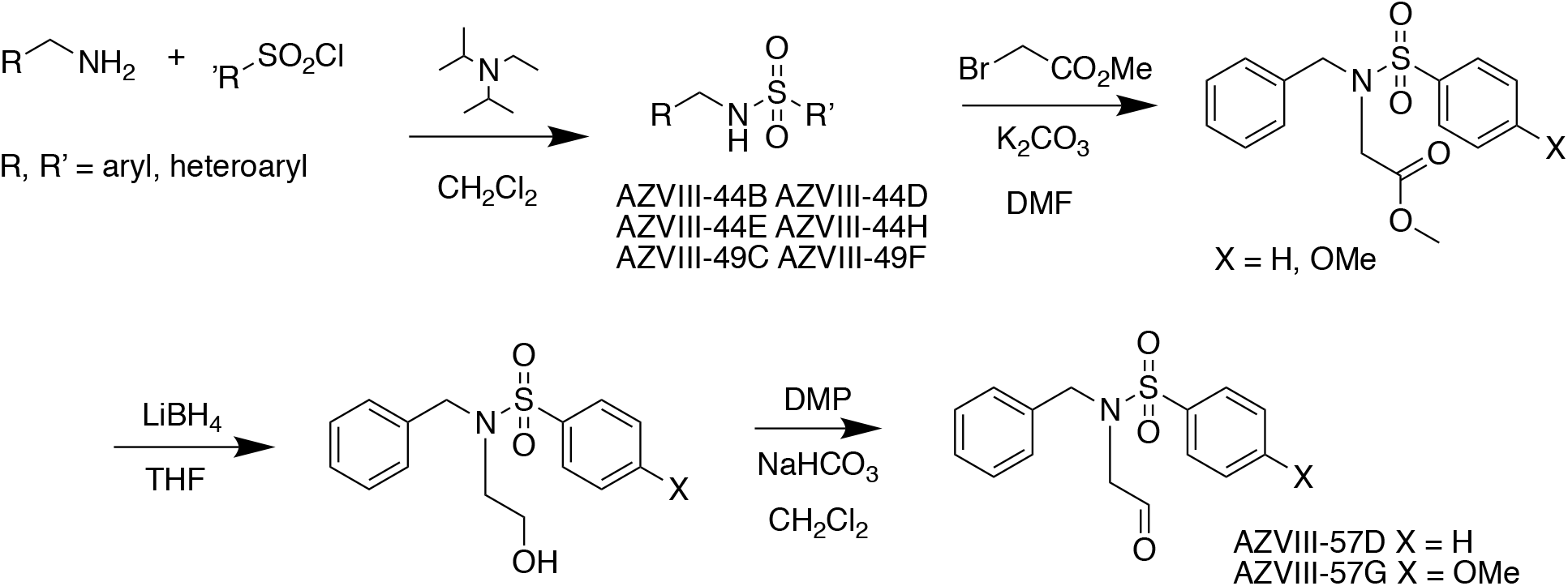

GC373 was synthesized from GC376 by converting the bisulfite group to an aldehyde group by treatment with aqueous sodium bicarbonate. To a solution of 5.31 mg of GC376 in 200 μL H_2_O, 2 μL of saturated NaHCO_3_ was added. The cloudy mixture was extracted with CH_2_Cl_2_, dried over Na_2_SO_4_, and concentrated in vacuo to give GC373 as a colorless oil (4.0 mg).

## Data availability

All reagents generated in this study are without restriction. Plasmids generated in this study will be deposited to Addgene. Source data for all figures are provided with this manuscript online. All statistics were performed using Prism v.8.4.2.

## Acknowledgements

Compound 11a was synthesized and generously provided by Shi-Xian Deng, Columbia University. This work was supported by a grant from the Jack Ma Foundation to D.D.H. and A.C. and by grants from Columbia Technology Ventures and the Columbia Translational Therapeutics (TRx) program to B.R.S. Further support for these studies comes from a pilot grant award from the Herbert Irving Comprehensive Cancer Center in partnership with the Irving Institute for Clinical and Translational Research at Columbia University to H.Y. and A.C. A.C. is also supported by a Career Awards for Medical Scientists from the Burroughs Wellcome Fund. S.J.R is supported by NIH grant F31NS111851. S.I. is supported by NIH grant T32AI106711.

## Author contributions

S.J.R., S.I, and A.C conceived the project. S.J.R., S.I., B.R.S., D.D.H., and A.C. planned and designed the experiments. S.J.R. performed crystal violet-based assays. S.J.R. and S.K. performed the HEK293-EYFP based assays. S.J.R., S.I., and S.J.H. cloned plasmids. A.Z. and H.L. synthesized compounds and provided compound structure information for synthesized compounds. N.E.S.T. and T.R. synthesized compounds NT-1-21, NT-1-24, and NT-1-32. M.S.N. and Y.H. conducted the live virus assays. S.K. and H.Y. performed imaging and created the HEK293-EYFP cell line. S.J.R. and S.M. conducted data analysis. S.J.R, S.I., S.J.H, and A.C. wrote the manuscript with input from all authors.

## Competing interests

S.I., H.L., A.Z., B.R.S., A.C., and D.D.H. are inventors on a patent application submitted based on some of the molecules described in this work. B.R.S. is an inventor on additional patents and patent applications related to small molecule therapeutics, and co-founded and serves as a consultant to Inzen Therapeutics and Nevrox Limited.

## References

1. Wu, F. et al. A new coronavirus associated with human respiratory disease in China. Nature 579, 265–269 (2020).

2. Zhou, P. et al. A pneumonia outbreak associated with a new coronavirus of probable bat origin. Nature 579, 270–273 (2020).

3. Pillaiyar, T., Manickam, M., Namasivayam, V., Hayashi, Y. & Jung, S.-H. An Overview of Severe Acute Respiratory Syndrome–Coronavirus (SARS-CoV) 3CL Protease Inhibitors: Peptidomimetics and Small Molecule Chemotherapy. J. Med. Chem. 59, 6595–6628 (2016).

4. de Wit, E., van Doremalen, N., Falzarano, D. & Munster, V. J. SARS and MERS: recent insights into emerging coronaviruses. Nat. Rev. Microbiol. 14, 523–534 (2016).

5. Huang, C., Wei, P., Fan, K., Liu, Y. & Lai, L. 3C-like Proteinase from SARS Coronavirus Catalyzes Substrate Hydrolysis by a General Base Mechanism ^†^. Biochemistry 43, 4568–4574 (2004).

6. Berry, M., Fielding, B. & Gamieldien, J. Human coronavirus OC43 3CL protease and the potential of ML188 as a broad-spectrum lead compound: Homology modelling and molecular dynamic studies. BMC Struct. Biol. 15, (2015).

7. Blanco, R., Carrasco, L. & Ventoso, I. Cell Killing by HIV-1 Protease. J. Biol. Chem. 278, 1086–1093 (2003).

8. Li, H. et al. Zika Virus Protease Cleavage of Host Protein Septin-2 Mediates Mitotic Defects in Neural Progenitors. Neuron 101, 1089–1098.e4 (2019).

9. Chau, D. H. W. et al. Coxsackievirus B3 proteases 2A and 3C induce apoptotic cell death through mitochondrial injury and cleavage of eIF4GI but not DAP5/p97/NAT1. Apoptosis 12, 513–524 (2007).

10. M’Barek, N. B., Audoly, G., Raoult, D. & Gluschankof, P. HIV-2 Protease resistance defined in yeast cells. Retrovirology 3, 58 (2006).

11. Barco, A., Feduchi, E. & Carrasco, L. Poliovirus Protease 3Cpro Kills Cells by Apoptosis. Virology 266, 352–360 (2000).

12. Li, M.-L. et al. The 3C protease activity of enterovirus 71 induces human neural cell apoptosis. Virology 293, 386–395 (2002).

13. Jin, Z. et al. Structure of Mpro from SARS-CoV-2 and discovery of its inhibitors. Nature 582, 289–293 (2020).

14. Zhu, W. et al. Identification of SARS-CoV-2 3CL Protease Inhibitors by a Quantitative High-throughput Screening. bioRxiv 2020.07.17.207019 (2020) doi:10.1101/2020.07.17.207019.

15. CDC. Information for Laboratories about Coronavirus (COVID-19). Centers for Disease Control and Prevention https://www.cdc.gov/coronavirus/2019-ncov/lab/lab-biosafety-guidelines.html (2020).

16. Pyrc, K. et al. Culturing the Unculturable: Human Coronavirus HKU1 Infects, Replicates, and Produces Progeny Virions in Human Ciliated Airway Epithelial Cell Cultures. J. Virol. 84, 11255–11263 (2010).

17. Sacco, M. D. et al. Structure and inhibition of the SARS-CoV-2 main protease reveals strategy for developing dual inhibitors against Mpro and cathepsin L. bioRxiv 2020.07.27.223727 (2020) doi:10.1101/2020.07.27.223727.

18. Sies, H. & Parnham, M. J. Potential therapeutic use of ebselen for COVID-19 and other respiratory viral infections. Free Radic. Biol. Med. 156, 107–112 (2020).

19. Feoktistova, M., Geserick, P. & Leverkus, M. Crystal Violet Assay for Determining Viability of Cultured Cells. Cold Spring Harb. Protoc. 2016, pdb.prot087379 (2016).

20. Ma, C. et al. Boceprevir, GC-376, and calpain inhibitors II, XII inhibit SARS-CoV-2 viral replication by targeting the viral main protease. Cell Res. 1–15 (2020) doi:10.1038/s41422-020-0356-z.

21. Vuong, W. et al. Feline coronavirus drug inhibits the main protease of SARS-CoV-2 and blocks virus replication. bioRxiv 2020.05.03.073080 (2020) doi:10.1101/2020.05.03.073080.

22. Froggatt, H. M., Heaton, B. E. & Heaton, N. S. Development of a fluorescence based, high-throughput SARS-CoV-2 3CL^pro^ reporter assay. bioRxiv 2020.06.24.169565 (2020) doi:10.1101/2020.06.24.169565.

23. Luan, X. et al. Structure Basis for Inhibition of SARS-CoV-2 by the Feline Drug GC376. bioRxiv 2020.06.07.138677 (2020) doi:10.1101/2020.06.07.138677.

24. Iketani, S. et al. Lead compounds for the development of SARS-CoV-2 3CL protease inhibitors. bioRxiv 2020.08.03.235291 (2020) doi:10.1101/2020.08.03.235291.

25. Yang, S. et al. Synthesis, Crystal Structure, Structure−Activity Relationships, and Antiviral Activity of a Potent SARS Coronavirus 3CL Protease Inhibitor. J. Med. Chem. 49, 4971–4980 (2006).

26. Dai, W. et al. Structure-based design of antiviral drug candidates targeting the SARS-CoV-2 main protease. Science 368, 1331–1335 (2020).

27. Ghosh, A. K. et al. Design, synthesis and antiviral efficacy of a series of potent chloropyridyl ester-derived SARS-CoV 3CLpro inhibitors. Bioorg. Med. Chem. Lett. 18, 5684–5688 (2008).

28. Węglarz-Tomczak, E., Tomczak, J. M., Talma, M. & Brul, S. Ebselen as a highly active inhibitor of PL^Pro^CoV2. bioRxiv 2020.05.17.100768 (2020) doi:10.1101/2020.05.17.100768.

29. Seale, L. A., Torres, D. J., Berry, M. J. & Pitts, M. W. A role for selenium-dependent GPX1 in SARS-CoV-2 virulence. Am. J. Clin. Nutr. doi:10.1093/ajcn/nqaa177.

30. Kim, Y. et al. Reversal of the Progression of Fatal Coronavirus Infection in Cats by a Broad-Spectrum Coronavirus Protease Inhibitor. PLOS Pathog. 12, e1005531 (2016).

31. Zhang, J.-H., Chung, T. D. Y. & Oldenburg, K. R. A Simple Statistical Parameter for Use in Evaluation and Validation of High Throughput Screening Assays. J. Biomol. Screen. 4, 67–73 (1999).

32. Birmingham, A. et al. Statistical Methods for Analysis of High-Throughput RNA Interference Screens. Nat. Methods 6, 569–575 (2009).

33. Kim, Y. et al. Broad-Spectrum Antivirals against 3C or 3C-Like Proteases of Picornaviruses, Noroviruses, and Coronaviruses. J. Virol. 86, 11754–11762 (2012).

34. Blanchard, J. E. et al. High-throughput screening identifies inhibitors of the SARS coronavirus main proteinase. Chem. Biol. 11, 1445–1453 (2004).

35. Klotz, L. C. & Sylvester, E. J. The Consequences of a Lab Escape of a Potential Pandemic Pathogen. Front. Public Health 2, (2014).

36. Coelho, A. C. & García Díez, J. Biological Risks and Laboratory-Acquired Infections: A Reality That Cannot be Ignored in Health Biotechnology. Front. Bioeng. Biotechnol. 3, (2015).

37. Tan, C. W. et al. A SARS-CoV-2 surrogate virus neutralization test based on antibody-mediated blockage of ACE2–spike protein–protein interaction. Nat. Biotechnol. (2020) doi:10.1038/s41587-020-0631-z.

38. Case, J. B. et al. Neutralizing Antibody and Soluble ACE2 Inhibition of a Replication-Competent VSV-SARS-CoV-2 and a Clinical Isolate of SARS-CoV-2. Cell Host Microbe doi:10.1016/j.chom.2020.06.021.

39. Dieterle, M. E. et al. A Replication-Competent Vesicular Stomatitis Virus for Studies of SARS-CoV-2 Spike-Mediated Cell Entry and Its Inhibition. Cell Host Microbe doi:10.1016/j.chom.2020.06.020.

40. Iglewicz, B. & Hoaglin, D. C. How to detect and handle outliers. (ASQC Quality Press, 1993).

41. Metzger, A., Melzig, L., Despotopoulou, C. & Knochel, P. Pd-Catalyzed Cross-Coupling of Functionalized Organozinc Reagents with Thiomethyl-Substituted Heterocycles. Org. Lett. 11, 4228–4231 (2009).

42. Zambaldo, C. et al. 2-Sulfonylpyridines as Tunable, Cysteine-Reactive Electrophiles. J. Am. Chem. Soc. 142, 8972–8979 (2020).

43. Mou, K. et al. Novel CADD-based peptidyl vinyl ester derivatives as potential proteasome inhibitors. Bioorg. Med. Chem. Lett. 18, 2198–2202 (2008).

44. Amblard, F. et al. Synthesis and antiviral evaluation of novel peptidomimetics as norovirus protease inhibitors. Bioorg. Med. Chem. Lett. 28, 2165–2170 (2018).

