## Supplementary Data for "A simplified cell-based assay to identify coronavirus 3CL protease inhibitors"

**Supplementary Fig. 1. Dose response experiments with SARS-CoV-2 3CLpro and GC376 are robust to variable assay parameters. a-d**. Repetition of assay with variable levels of cell seeding into drug conditions result in similar $EC_{50}$ value predictions. **e-h**. Repetition of assay with variable levels of plasmid transfection results in similar $EC_{50}$ value predictions. $EC_{50}$ values are displayed as best-fit value alongside 95% confidence interval. Data are shown as mean ± s.d. for four technical replicates.

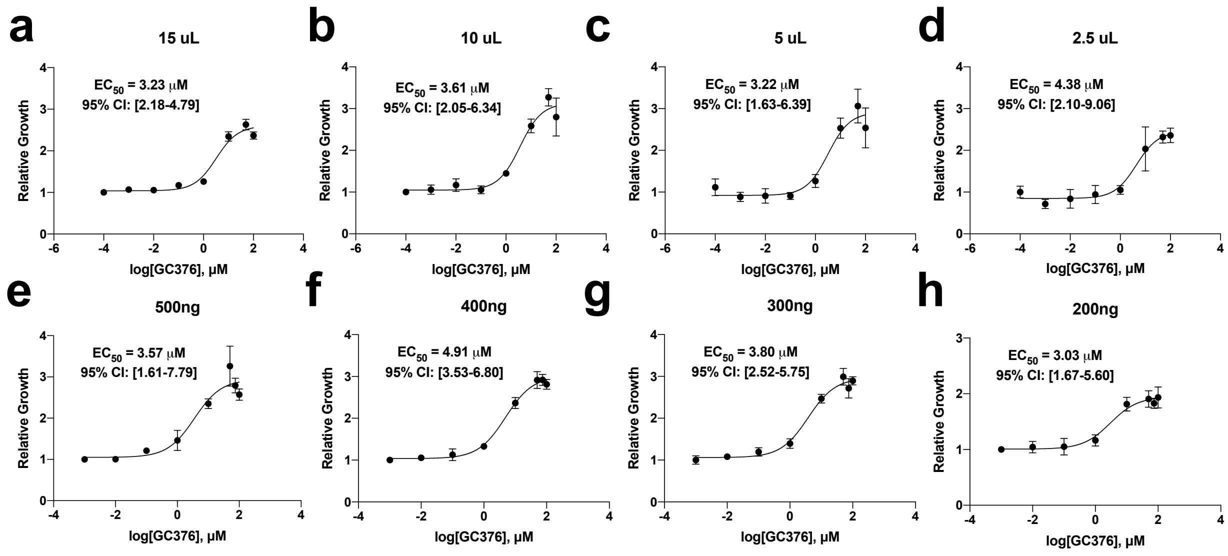

**Supplementary Fig. 2. Dose response experiments with SARS-CoV-2 3CLpro can be determined with imaging of EYFP labeled HEK293 cells. a-b**. The activity of GC376 and compound 4 can be detected using imaging rather than crystal violet staining. $EC_{50}$ values are displayed as best-fit value alongside 95% confidence interval. Data are shown as mean ± s.d. for four technical replicates.

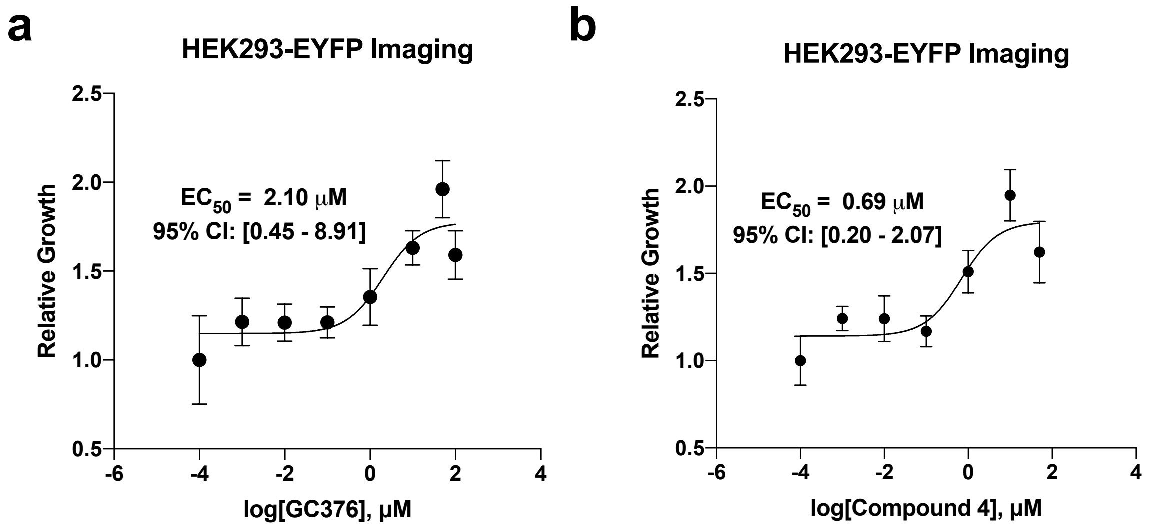

**Supplementary Figure 3. Live virus testing of compound 11a.** The $EC_{50}$ value is displayed as best-fit value alongside 95% confidence interval. Data are shown as mean ± s.d. for three technical replicates.

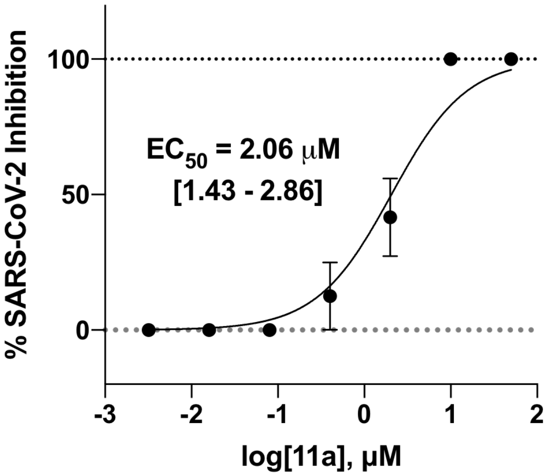

**Supplementary Fig. 4. Compounds with activity against the SARS-CoV-2 3CLpro *in vitro* that are not efficacious against the SARS-CoV-2 live virus do not show activity in the transfection-based assay. a**. DMSO was tested at concentrations up to 1%, the maximal concentration used to deliver compounds in this study, and does not show toxicity to HEK293T cells or protease inhibitory activity against the 3CLpro. **b-e**. Other compounds with reported activity against purified SARS-CoV-2 3CLpro but not against live virus. **f**. Ebselen, a compound with efficacy against the SARS-CoV-2 live virus does not rescue 3CLpro induced cytotoxicity within the transfection-based assay. $EC_{50}$ values are displayed as best-fit value alongside 95% confidence interval. $CC_{50}$ values are displayed as best-fit value. Data are shown as mean ± s.d. for three or four technical replicates.

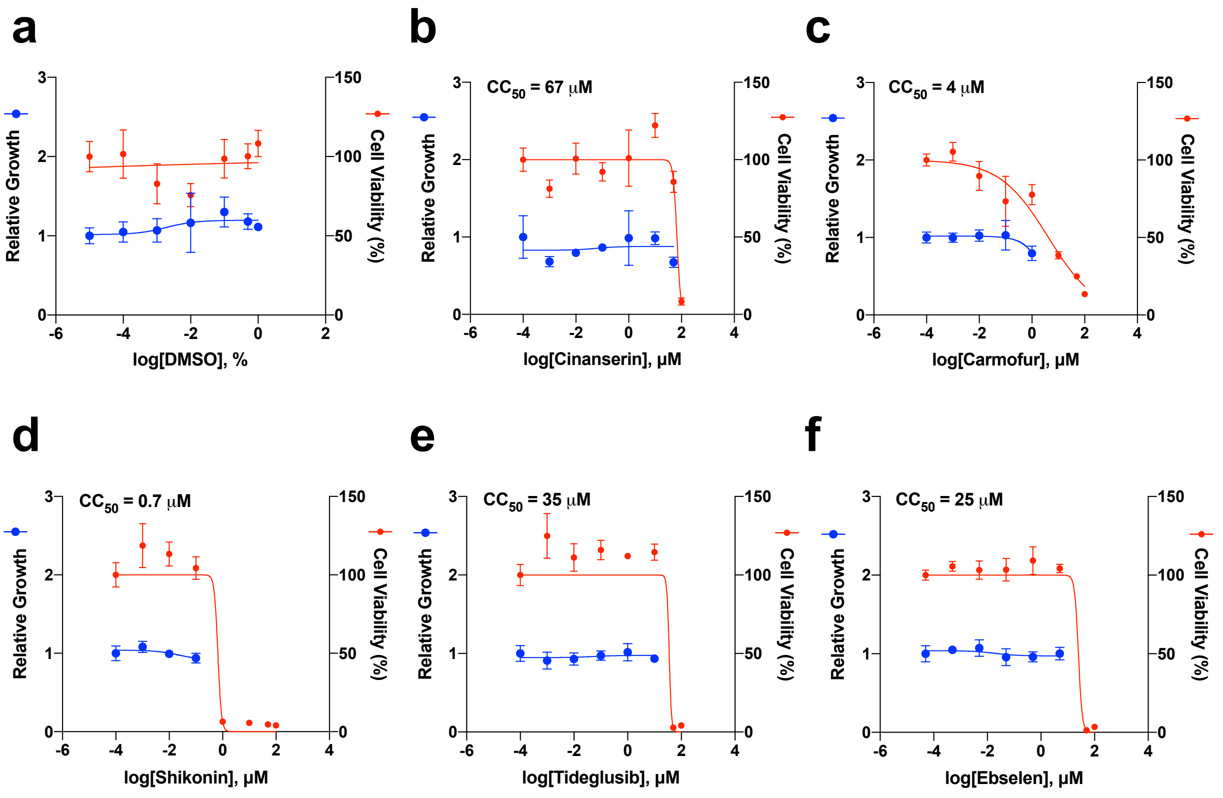

**Supplementary Fig. 5. A number of other 3CLpro enzymes from different coronavirus species also show activity-dependent cytotoxicity. a**. A phylogenetic tree of the six coronaviruses tested in this study, generated using NCBI Virus with the sequences of the ORF1ab polyprotein. The genera of each virus are shown along with the amino acid sequence similarity to the SARS-CoV-2 3CLpro calculated using the Protein BLAST tool from NCBI. **b**. Quantification of cytotoxicity upon expression of active or inactivated 3CLpro enzymes from SARS-CoV, MERS-CoV, Bat-CoV-HKU9, HCoV-NL63, and IBV in 293T cells. Data are shown as mean ± s.d. for four technical replicates.

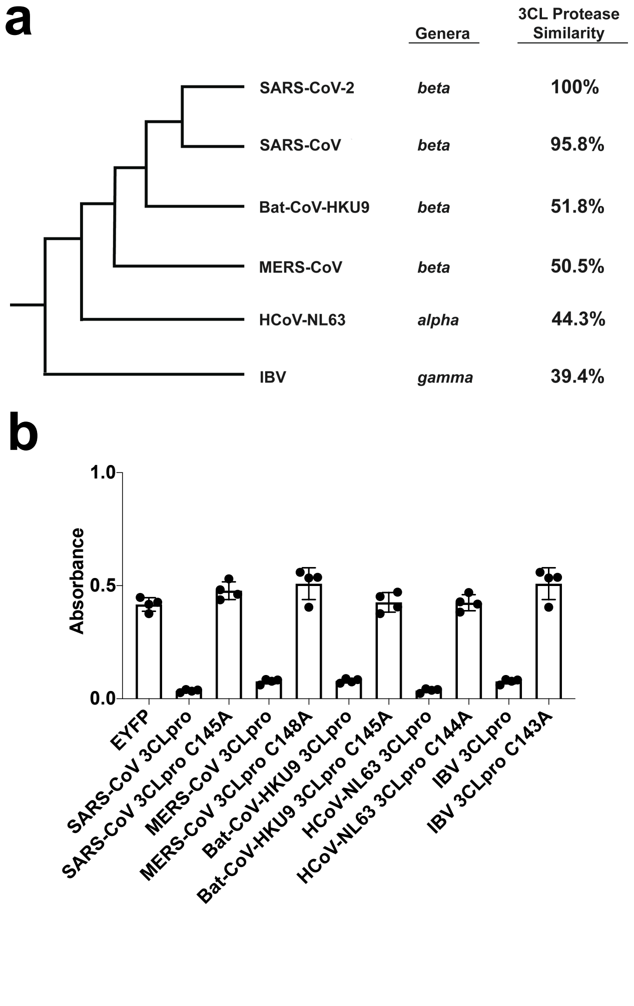

**Supplementary Fig. 6 Optimization of screening parameters and resulting Z-factors between positive and negative control wells.** Z-factor, a measure of assay quality was determined for two positive control compounds, GC376 at 50 μM and compound 4 at 20 μM, at different time points to identify optimal screening conditions for compounds against the SARS-CoV-2-3CLpro. The DMSO condition was conducted with 21 technical replicates randomly positioned across a 96-well plate while positive control compounds were tested at three or five technical replicates. FC = Fold Change. Data are shown as mean ± s.d. for specified technical replicates.

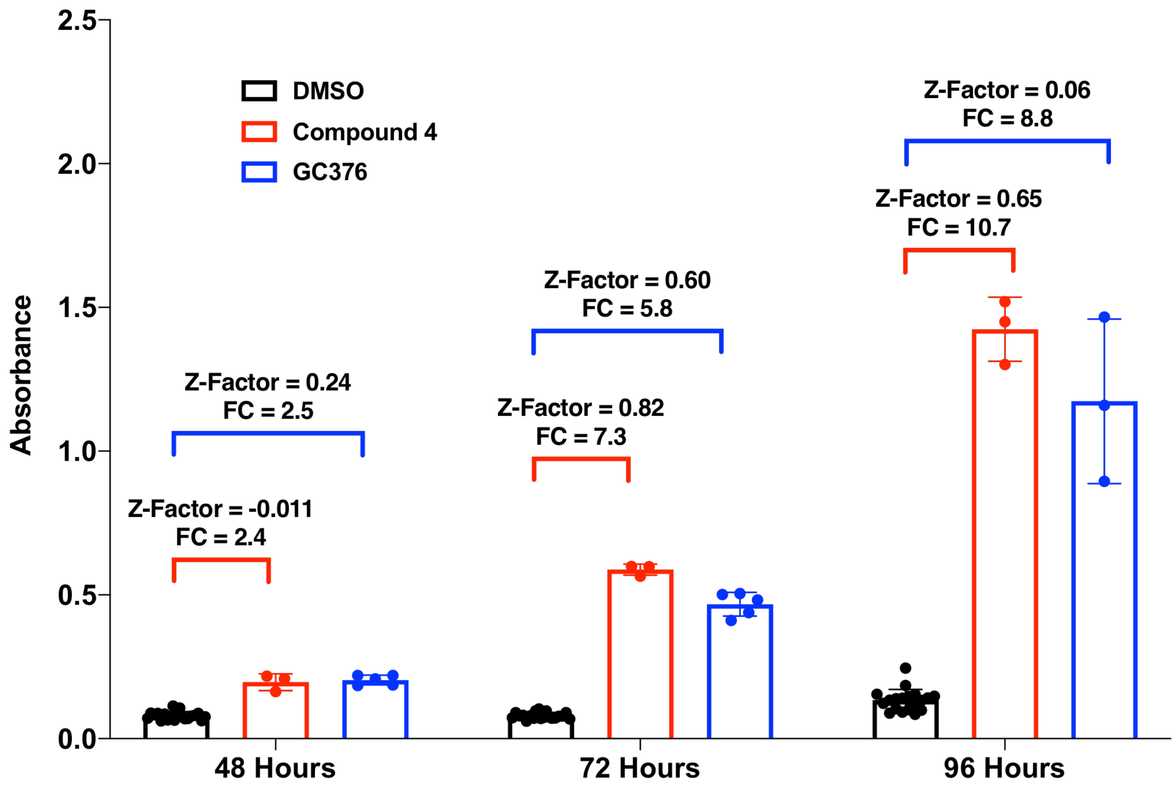

**Supplementary Fig. 7. GC373 demonstrates activity against the SARS-CoV-2 3CLpro and against the SARS-CoV-2 live virus. a**. Dose-response profiling and cytotoxicity determination using the transfection-based assay of GC373 against the SARS-CoV-2 3CLpro. **b**. Live virus testing of GC373 against SARS-CoV-2. $EC_{50}$ values are displayed as best-fit value alongside 95% confidence interval. The live virus assay was conducted with two biological replicates, each with three technical replicates and the $EC_{50}$ value was derived from all replicates. $CC_{50}$ values are displayed as best-fit value. Data are shown as mean ± s.d. for three or four technical replicates.

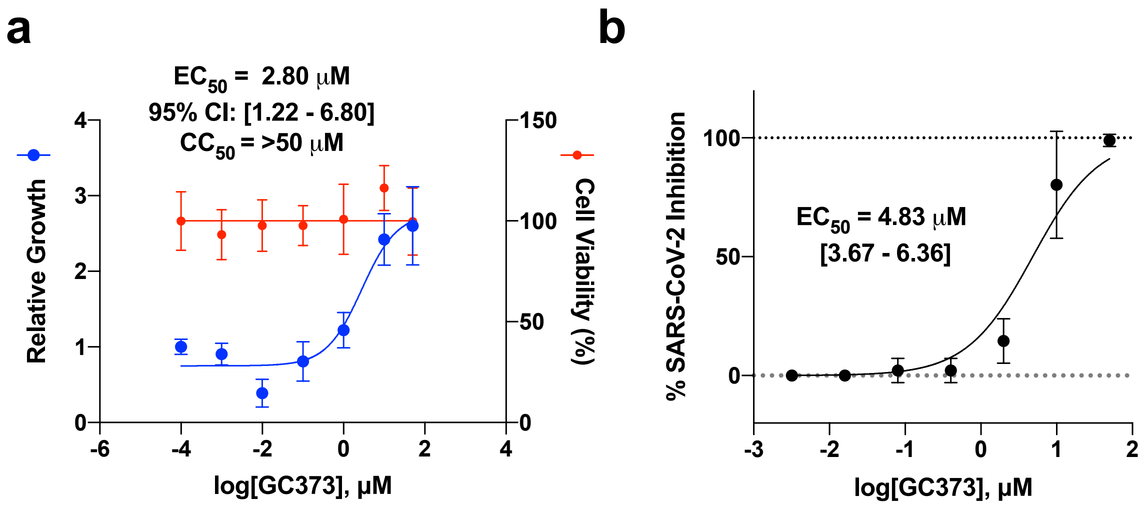

**Supplementary Fig. 8. The activity of GRL-0496 and GC373 show variable efficacy and potency against the coronavirus 3CL proteases from SARS-CoV, MERS-CoV, Bat-CoV-HKU9, HCoV-NL63, and IBV.** $EC_{50}$ values are displayed as best-fit value alongside 95% confidence interval. $CC_{50}$ values are displayed as best-fit value. Data are shown as mean ± s.d. for four technical replicates.

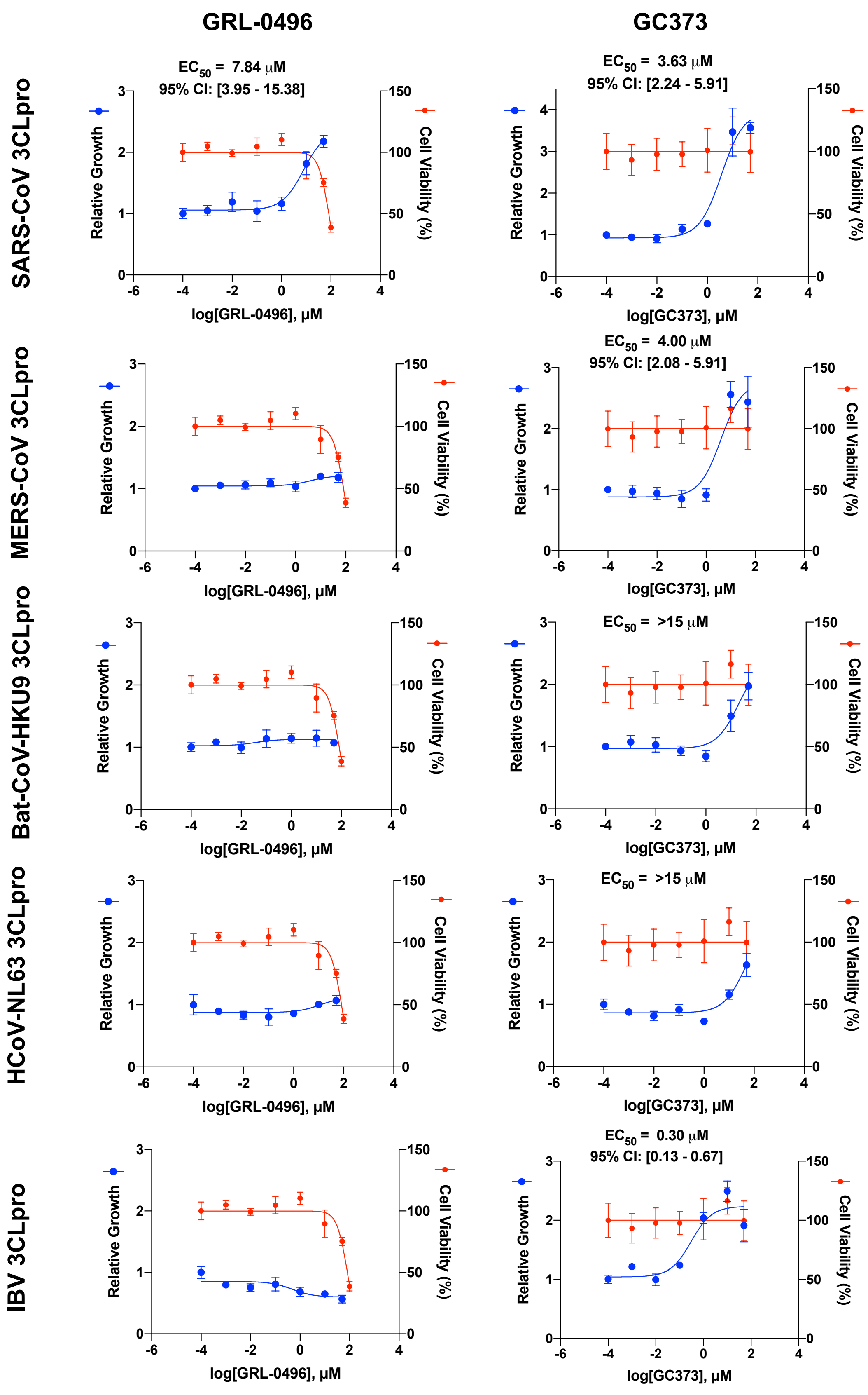
GRL-0496
GC373
SARS-CoV 3CLpro
MERS-CoV 3CLpro
Bat-CoV-HKU9 3CLpro
HCoV-NL63 3CLpro
IBV 3CLpro
EC50 = 7.84 μM
95% CI: [3.95 - 15.38]
EC50 = 3.63 μM
95% CI: [2.24 - 5.91]
EC50 = 4.00 μM
95% CI: [2.08 - 5.91]
EC50 = >15 μM
EC50 = >15 μM
EC50 = 0.30 μM
95% CI: [0.13 - 0.67]
Relative Growth
Cell Viability (%)
log[GRL-0496], μM
log[GC373], μM

**Supplementary Table 1. Selectivity Index (SI) for compounds tested with the transfection-based assay in this study.**

| Protease | Drug | $EC_{50}$ (μM) | $CC_{50}$ (μM) | SI ($CC_{50}$ /$EC_{50}$) |
|---|---|---|---|---|
| SARS-CoV-2 3CLpro | GC376 | 3.3 | >100 | >30.3 |
| | compound 4 | 0.98 | 81 | 82.7 |
| | 11a | 6.89 | 48 | 7 |
| | GRL-0496 | 5.05 | 81 | 16 |
| | GC373 | 2.8 | >50 | >17.9 |
| SARS-CoV 3CLpro | GC376 | 5.83 | >100 | >17.2 |
| | compound 4 | 3.17 | 81 | 25.6 |
| | 11a | 5.38 | 48 | 8.9 |
| | GRL-0496 | 7.84 | 81 | 10.3 |
| | GC373 | 3.63 | >50 | >13.8 |
| MERS-CoV 3CLpro | GC376 | 7.44 | >100 | >13.4 |
| | compound 4 | 1.4 | 81 | 57.9 |
| | GC373 | 4 | >50 | >12.5 |
| Bat-CoV-HKU9 3CLpro | GC376 | 11.07 | >100 | >9.0 |
| | compound 4 | 4.11 | 81 | 19.7 |
| HCoV-NL63 3CLpro | compound 4 | 4.92 | 81 | 16.5 |
| IBV 3CLpro | GC376 | 0.58 | >100 | >172 |
| | compound 4 | 0.058 | 81 | 1396.5 |
| | GC373 | 0.3 | >50 | >166.6 |

**Supplementary Table 2. Compounds screened for activity against the SARS-CoV-2 3CLpro.**

| Absorbance | Well | Drug | Plate | Model | z-score |
|---|---|---|---|---|---|
| 0.07156 | A02 | Omarigliptin | Plate1 | SARS-CoV-2 3CLpro | 0.07462451 |
| 0.03996 | A03 | Apoptosis Activator 2 | Plate1 | SARS-CoV-2 3CLpro | -3.5532747 |
| 0.06801 | A04 | Picolamine | Plate1 | SARS-CoV-2 3CLpro | -0.3329401 |
| 0.07551 | A05 | Muscone | Plate1 | SARS-CoV-2 3CLpro | 0.52811191 |
| 0.08101 | A06 | 2-Aminoethanethiol | Plate1 | SARS-CoV-2 3CLpro | 1.15955007 |
| 0.07101 | A07 | Dexibuprofen | Plate1 | SARS-CoV-2 3CLpro | 0.01148069 |
| 0.07496 | A08 | Glucosamine | Plate1 | SARS-CoV-2 3CLpro | 0.4649681 |
| 0.07141 | A09 | Gabexate mesylate | Plate1 | SARS-CoV-2 3CLpro | 0.05740347 |
| 0.09891 | A10 | Zalcitabine | Plate1 | SARS-CoV-2 3CLpro | 3.21459426 |
| 0.09016 | A11 | Amiloride hydrochloride | Plate1 | SARS-CoV-2 3CLpro | 2.21003355 |
| 0.06776 | B02 | Saxagliptin hydrate | Plate1 | SARS-CoV-2 3CLpro | -0.3616419 |
| 0.07901 | B03 | Linagliptin | Plate1 | SARS-CoV-2 3CLpro | 0.9299362 |
| 0.07831 | B04 | Sitagliptin | Plate1 | SARS-CoV-2 3CLpro | 0.84957134 |
| 0.08006 | B05 | Hexylresorcinol | Plate1 | SARS-CoV-2 3CLpro | 1.05048348 |
| 0.06906 | B06 | Arbutin | Plate1 | SARS-CoV-2 3CLpro | -0.2123928 |
| 0.03171 | B07 | Diminazene Aceturate | Plate1 | SARS-CoV-2 3CLpro | -4.500432 |
| 0.06266 | B08 | 3-Pyridylacetic acid hydrochloride | Plate1 | SARS-CoV-2 3CLpro | -0.9471572 |
| 0.05386 | B09 | Racecadotril | Plate1 | SARS-CoV-2 3CLpro | -1.9574583 |
| 0.06456 | B10 | Mizoribine | Plate1 | SARS-CoV-2 3CLpro | -0.7290241 |
| 0.08056 | B11 | Sodium etidronate | Plate1 | SARS-CoV-2 3CLpro | 1.10788695 |
| 0.07191 | C02 | MAC-5576 | Plate1 | SARS-CoV-2 3CLpro | 0.11480694 |
| 0.07666 | C03 | DMSO | Plate1 | SARS-CoV-2 3CLpro | 0.66013989 |
| 0.07206 | C04 | DMSO | Plate1 | SARS-CoV-2 3CLpro | 0.13202798 |
| 0.07451 | C05 | DMSO | Plate1 | SARS-CoV-2 3CLpro | 0.41330498 |
| 0.07591 | C06 | BTB07404 | Plate1 | SARS-CoV-2 3CLpro | 0.57403469 |
| 0.06836 | C07 | DMSO | Plate1 | SARS-CoV-2 3CLpro | -0.2927577 |
| 0.06826 | C08 | DMSO | Plate1 | SARS-CoV-2 3CLpro | -0.3042384 |
| 0.07401 | C09 | Myrecetin | Plate1 | SARS-CoV-2 3CLpro | 0.35590151 |
| 0.06881 | C10 | DMSO | Plate1 | SARS-CoV-2 3CLpro | -0.2410946 |
| 0.08486 | C11 | Tipranivir | Plate1 | SARS-CoV-2 3CLpro | 1.60155678 |
| 0.06106 | D02 | DMSO | Plate1 | SARS-CoV-2 3CLpro | -1.1308483 |
| 0.06566 | D03 | BTB07408 | Plate1 | SARS-CoV-2 3CLpro | -0.6027364 |
| 0.36221 | D04 | GC376 | Plate1 | SARS-CoV-2 3CLpro | 33.443261 |
| 0.06851 | D05 | MAC-8120 | Plate1 | SARS-CoV-2 3CLpro | -0.2755367 |
| 0.06866 | D06 | DMSO | Plate1 | SARS-CoV-2 3CLpro | -0.2583156 |
| 0.07186 | D07 | MWP00332 | Plate1 | SARS-CoV-2 3CLpro | 0.10906659 |

| | | | | | |
|---|---|---|---|---|---|
| 0.07301 | D08 | DMSO | Plate1 | SARS-CoV-2 3CLpro | 0.24109457 |
| 0.07556 | D09 | DMSO | Plate1 | SARS-CoV-2 3CLpro | 0.53385226 |
| 0.07011 | D10 | Rupintrivir | Plate1 | SARS-CoV-2 3CLpro | -0.0918456 |
| 0.06876 | D11 | DMSO | Plate1 | SARS-CoV-2 3CLpro | -0.2468349 |
| 0.06221 | E02 | DMSO | Plate1 | SARS-CoV-2 3CLpro | -0.9988204 |
| 0.06836 | E03 | DMSO | Plate1 | SARS-CoV-2 3CLpro | -0.2927577 |
| 0.06486 | E04 | BTB07417 | Plate1 | SARS-CoV-2 3CLpro | -0.694582 |
| 0.07096 | E05 | DMSO | Plate1 | SARS-CoV-2 3CLpro | 0.00574035 |
| 0.06971 | E06 | AZVIII-49F | Plate1 | SARS-CoV-2 3CLpro | -0.1377683 |
| 0.06231 | E07 | DMSO | Plate1 | SARS-CoV-2 3CLpro | -0.9873397 |
| 0.07276 | E08 | MWP00508 | Plate1 | SARS-CoV-2 3CLpro | 0.21239284 |
| 0.06846 | E09 | DMSO | Plate1 | SARS-CoV-2 3CLpro | -0.281277 |
| 0.06741 | E10 | DMSO | Plate1 | SARS-CoV-2 3CLpro | -0.4018243 |
| 0.07371 | E11 | MWP00333 | Plate1 | SARS-CoV-2 3CLpro | 0.32145943 |
| 0.04971 | F02 | Grazoprevir | Plate1 | SARS-CoV-2 3CLpro | -2.4339071 |
| 0.06991 | F03 | AZVIII-57D | Plate1 | SARS-CoV-2 3CLpro | -0.1148069 |
| 0.06831 | F04 | DMSO | Plate1 | SARS-CoV-2 3CLpro | -0.298498 |
| 0.06911 | F05 | BTB07407 | Plate1 | SARS-CoV-2 3CLpro | -0.2066525 |
| 0.07516 | F06 | DMSO | Plate1 | SARS-CoV-2 3CLpro | 0.48792949 |
| 0.47481 | F07 | CMPD18-20 | Plate1 | SARS-CoV-2 3CLpro | 46.3705222 |
| 0.07086 | F08 | DMSO | Plate1 | SARS-CoV-2 3CLpro | -0.0057403 |
| 0.18266 | F09 | GRL-0496 | Plate1 | SARS-CoV-2 3CLpro | 12.8296753 |
| 0.05571 | F10 | Saquinavir | Plate1 | SARS-CoV-2 3CLpro | -1.7450655 |
| 0.06161 | F11 | DMSO | Plate1 | SARS-CoV-2 3CLpro | -1.0677045 |
| 0.04941 | G02 | Monobenzone | Plate1 | SARS-CoV-2 3CLpro | -2.4683492 |
| 0.08226 | G03 | Limonin | Plate1 | SARS-CoV-2 3CLpro | 1.30305874 |
| 0.02596 | G04 | Betulinic acid | Plate1 | SARS-CoV-2 3CLpro | -5.1605719 |
| 0.07796 | G05 | PMSF | Plate1 | SARS-CoV-2 3CLpro | 0.80938891 |
| 0.08286 | G06 | Fenofibric acid | Plate1 | SARS-CoV-2 3CLpro | 1.37194291 |
| 0.07241 | G07 | Ramelteon | Plate1 | SARS-CoV-2 3CLpro | 0.17221041 |
| 0.05871 | G08 | Ritonavir | Plate1 | SARS-CoV-2 3CLpro | -1.4006446 |
| 0.08216 | G09 | Alogliptin Benzoate | Plate1 | SARS-CoV-2 3CLpro | 1.29157805 |
| 0.00986 | G10 | Bortezomib | Plate1 | SARS-CoV-2 3CLpro | -7.0089636 |
| 0.08641 | G11 | Acetohydroxamic acid | Plate1 | SARS-CoV-2 3CLpro | 1.77950754 |
| 0.07081 | H02 | 8 | Plate1 | SARS-CoV-2 3CLpro | -0.0114807 |
| 0.05576 | H03 | Lopinavir | Plate1 | SARS-CoV-2 3CLpro | -1.7393251 |
| 0.07581 | H04 | Penciclovir | Plate1 | SARS-CoV-2 3CLpro | 0.562554 |
| 0.01181 | H05 | AOB2796 | Plate1 | SARS-CoV-2 3CLpro | -6.78509 |
| 0.08421 | H06 | Maribavir | Plate1 | SARS-CoV-2 3CLpro | 1.52693227 |

| | | | | | |
|---|---|---|---|---|---|
| 0.08156 | H07 | Trelagliptin succinate | Plate1 | SARS-CoV-2 3CLpro | 1.22269389 |
| 0.01741 | H08 | MLN9708 | Plate1 | SARS-CoV-2 3CLpro | -6.1421712 |
| 0.07691 | H09 | SC514 | Plate1 | SARS-CoV-2 3CLpro | 0.68884163 |
| 0.01011 | H10 | Ixazomib | Plate1 | SARS-CoV-2 3CLpro | -6.9802618 |
| 0.08826 | H11 | Raltegravir potassium | Plate1 | SARS-CoV-2 3CLpro | 1.99190037 |
| 0.082225 | A02 | PSI6206 | Plate2 | SARS-CoV-2 3CLpro | 0.48063475 |
| 0.069275 | A03 | Cilastatin | Plate2 | SARS-CoV-2 3CLpro | -0.3492612 |
| 0.068975 | A04 | Taxifolin | Plate2 | SARS-CoV-2 3CLpro | -0.3684866 |
| 0.084325 | A05 | Nafamostat mesylate | Plate2 | SARS-CoV-2 3CLpro | 0.61521247 |
| 0.047775 | A06 | Daclatasvir dihydrochloride | Plate2 | SARS-CoV-2 3CLpro | -1.7270809 |
| 0.094525 | A07 | Darunavir†Ethanolate | Plate2 | SARS-CoV-2 3CLpro | 1.26887573 |
| 0.106675 | A08 | Ilomastat | Plate2 | SARS-CoV-2 3CLpro | 2.04750402 |
| 0.049625 | A09 | Elvitegravir | Plate2 | SARS-CoV-2 3CLpro | -1.6085243 |
| 0.076575 | A10 | Dolutegravir sodium | Plate2 | SARS-CoV-2 3CLpro | 0.11855657 |
| 0.108725 | A11 | Astragaloside IV | Plate2 | SARS-CoV-2 3CLpro | 2.17887751 |
| 0.064175 | B02 | Arctigenin | Plate2 | SARS-CoV-2 3CLpro | -0.6760929 |
| 0.086575 | B03 | Stigmasterol | Plate2 | SARS-CoV-2 3CLpro | 0.7594029 |
| 0.054075 | B04 | Nobiletin | Plate2 | SARS-CoV-2 3CLpro | -1.3233477 |
| 0.005175 | B05 | Celastrol | Plate2 | SARS-CoV-2 3CLpro | -4.4570862 |
| 0.093275 | B06 | Glucosamine sulfate | Plate2 | SARS-CoV-2 3CLpro | 1.18876994 |
| 0.068025 | B07 | Picroside I | Plate2 | SARS-CoV-2 3CLpro | -0.429367 |
| 0.078775 | B08 | Alvelestat | Plate2 | SARS-CoV-2 3CLpro | 0.25954276 |
| 0.079225 | B09 | N-Ethylmaleimide | Plate2 | SARS-CoV-2 3CLpro | 0.28838085 |
| 0.068525 | B10 | DAPT | Plate2 | SARS-CoV-2 3CLpro | -0.3973247 |
| 0.103975 | B11 | Trelagliptin | Plate2 | SARS-CoV-2 3CLpro | 1.87447551 |
| 0.085725 | C02 | Fosamprenavir | Plate2 | SARS-CoV-2 3CLpro | 0.70493096 |
| 0.057525 | C03 | DMSO | Plate2 | SARS-CoV-2 3CLpro | -1.1022557 |
| 0.056875 | C04 | DMSO | Plate2 | SARS-CoV-2 3CLpro | -1.1439107 |
| 0.060275 | C05 | Indinavir | Plate2 | SARS-CoV-2 3CLpro | -0.9260229 |
| 0.081325 | C06 | DMSO | Plate2 | SARS-CoV-2 3CLpro | 0.42295858 |
| 0.085675 | C07 | DMSO | Plate2 | SARS-CoV-2 3CLpro | 0.70172673 |
| 0.726225 | C08 | CMPD18-20 | Plate2 | SARS-CoV-2 3CLpro | 41.7511382 |
| 0.067575 | C09 | DMSO | Plate2 | SARS-CoV-2 3CLpro | -0.4582051 |
| 0.117025 | C10 | SPB08384 | Plate2 | SARS-CoV-2 3CLpro | 2.71077996 |
| 0.123825 | C11 | DMSO | Plate2 | SARS-CoV-2 3CLpro | 3.14655547 |
| 0.085025 | D02 | DMSO | Plate2 | SARS-CoV-2 3CLpro | 0.66007172 |
| 0.486725 | D03 | CMPD18-10 | Plate2 | SARS-CoV-2 3CLpro | 26.4028686 8 |
| 0.074775 | D04 | DMSO | Plate2 | SARS-CoV-2 3CLpro | 0.00320423 |

| | | | | | |
|---|---|---|---|---|---|
| 0.075325 | D05 | DMSO | Plate2 | SARS-CoV-2 3CLpro | 0.03845078 |
| 0.088475 | D06 | Apigenin | Plate2 | SARS-CoV-2 3CLpro | 0.8811637 |
| 0.080225 | D07 | AZVIII-57G | Plate2 | SARS-CoV-2 3CLpro | 0.35246548 |
| 0.060775 | D08 | DMSO | Plate2 | SARS-CoV-2 3CLpro | -0.8939806 |
| 0.066125 | D09 | SPB06613 | Plate2 | SARS-CoV-2 3CLpro | -0.5511278 |
| 0.074225 | D10 | DMSO | Plate2 | SARS-CoV-2 3CLpro | -0.0320423 |
| 0.113225 | D11 | SPB06636 | Plate2 | SARS-CoV-2 3CLpro | 2.46725836 |
| 0.071675 | E02 | AZVIII-38 | Plate2 | SARS-CoV-2 3CLpro | -0.1954581 |
| 0.083175 | E03 | DMSO | Plate2 | SARS-CoV-2 3CLpro | 0.54151515 |
| 0.073475 | E04 | AZVIII-49C | Plate2 | SARS-CoV-2 3CLpro | -0.0801058 |
| 0.085225 | E05 | DMSO | Plate2 | SARS-CoV-2 3CLpro | 0.67288864 |
| 0.070925 | E06 | Quercetin | Plate2 | SARS-CoV-2 3CLpro | -0.2435216 |
| 0.078575 | E07 | DMSO | Plate2 | SARS-CoV-2 3CLpro | 0.24672584 |
| 0.073625 | E08 | SPB06591 | Plate2 | SARS-CoV-2 3CLpro | -0.0704931 |
| 0.076475 | E09 | DMSO | Plate2 | SARS-CoV-2 3CLpro | 0.11214811 |
| 0.077875 | E10 | SPB06593 | Plate2 | SARS-CoV-2 3CLpro | 0.20186659 |
| 0.117725 | E11 | DMSO | Plate2 | SARS-CoV-2 3CLpro | 2.75563921 |
| 0.074675 | F02 | DMSO | Plate2 | SARS-CoV-2 3CLpro | -0.0032042 |
| 0.060375 | F03 | DMSO | Plate2 | SARS-CoV-2 3CLpro | -0.9196145 |
| 0.077325 | F04 | DMSO | Plate2 | SARS-CoV-2 3CLpro | 0.16662005 |
| 0.078525 | F05 | Famotidine | Plate2 | SARS-CoV-2 3CLpro | 0.2435216 |
| 0.071325 | F06 | DMSO | Plate2 | SARS-CoV-2 3CLpro | -0.2178878 |
| 0.412325 | F07 | GC376 | Plate2 | SARS-CoV-2 3CLpro | 21.634972 |
| 0.070025 | F08 | DMSO | Plate2 | SARS-CoV-2 3CLpro | -0.3011978 |
| 0.068675 | F09 | DMSO | Plate2 | SARS-CoV-2 3CLpro | -0.387712 |
| 0.076275 | F10 | DMSO | Plate2 | SARS-CoV-2 3CLpro | 0.09933118 |
| 0.096375 | F11 | DMSO | Plate2 | SARS-CoV-2 3CLpro | 1.3874323 |
| 0.071175 | G02 | Z-VAD(OMe)-FMK | Plate2 | SARS-CoV-2 3CLpro | -0.2275004 |
| 0.076225 | G03 | Abietic Acid | Plate2 | SARS-CoV-2 3CLpro | 0.09612695 |
| 0.061225 | G04 | Atazanavir sulfate | Plate2 | SARS-CoV-2 3CLpro | -0.8651425 |
| 0.089475 | G05 | Abacavir | Plate2 | SARS-CoV-2 3CLpro | 0.94524833 |
| 0.056675 | G06 | Balicatib | Plate2 | SARS-CoV-2 3CLpro | -1.1567276 |
| 0.011625 | G07 | Carfilzomib | Plate2 | SARS-CoV-2 3CLpro | -4.0437403 |
| 0.054975 | G08 | Atazanavir | Plate2 | SARS-CoV-2 3CLpro | -1.2656715 |
| 0.068275 | G09 | Vildagliptin | Plate2 | SARS-CoV-2 3CLpro | -0.4133459 |
| 0.025775 | G10 | Dapivirine | Plate2 | SARS-CoV-2 3CLpro | -3.1369428 |
| 0.072375 | G11 | SB-3CT | Plate2 | SARS-CoV-2 3CLpro | -0.1505989 |
| 0.060825 | H02 | PD 151746 | Plate2 | SARS-CoV-2 3CLpro | -0.8907764 |
| 0.018775 | H03 | PAC1 | Plate2 | SARS-CoV-2 3CLpro | -3.5855352 |

| | | | | | |
|---|---|---|---|---|---|
| 0.073725 | H04 | Camostat mesilate | Plate2 | SARS-CoV-2 3CLpro | -0.0640846 |
| 0.073525 | H05 | Efavirenz | Plate2 | SARS-CoV-2 3CLpro | -0.0769016 |
| 0.088325 | H06 | Des(benzylpyridyl) Atazanavir | Plate2 | SARS-CoV-2 3CLpro | 0.87155101 |
| 0.073375 | H07 | LY2811376 | Plate2 | SARS-CoV-2 3CLpro | -0.0865143 |
| 0.013175 | H08 | FLI06 | Plate2 | SARS-CoV-2 3CLpro | -3.9444091 |
| 0.056425 | H09 | SRPIN340 | Plate2 | SARS-CoV-2 3CLpro | -1.1727488 |
| 0.096125 | H10 | NSC 405020 | Plate2 | SARS-CoV-2 3CLpro | 1.37141114 |
| 0.088525 | H11 | Leupeptin Hemisulfate | Plate2 | SARS-CoV-2 3CLpro | 0.88436793 |
| 0.01024 | A02 | Epoxomicin | Plate3 | SARS-CoV-2 3CLpro | -4.8596643 |
| 0.02879 | A03 | MG101 | Plate3 | SARS-CoV-2 3CLpro | -3.6239306 |
| 0.06649 | A04 | lavendustin C | Plate3 | SARS-CoV-2 3CLpro | -1.1124934 |
| 0.08099 | A05 | BMS707035 | Plate3 | SARS-CoV-2 3CLpro | -0.146556 |
| 0.05509 | A06 | Asunaprevir | Plate3 | SARS-CoV-2 3CLpro | -1.87192 |
| 0.07839 | A07 | Loxistatin Acid | Plate3 | SARS-CoV-2 3CLpro | -0.3197586 |
| 0.00484 | A08 | GK921 | Plate3 | SARS-CoV-2 3CLpro | -5.2193927 |
| 0.06729 | A09 | L-685,458 | Plate3 | SARS-CoV-2 3CLpro | -1.0592003 |
| 0.07469 | A10 | Tenofovir Disoproxil Fumarate | Plate3 | SARS-CoV-2 3CLpro | -0.5662392 |
| 0.02539 | A11 | GSK690693 | Plate3 | SARS-CoV-2 3CLpro | -3.8504263 |
| 0.03379 | B02 | Ledipasvir | Plate3 | SARS-CoV-2 3CLpro | -3.2908487 |
| 0.01554 | B03 | ONX0914 | Plate3 | SARS-CoV-2 3CLpro | -4.5065975 |
| 0.06614 | B04 | PI1840 | Plate3 | SARS-CoV-2 3CLpro | -1.1358091 |
| 0.08239 | B05 | (+)-Isocorydine hydrochloride | Plate3 | SARS-CoV-2 3CLpro | -0.0532931 |
| 0.09179 | B06 | UAMC 00039 dihydrochloride | Plate3 | SARS-CoV-2 3CLpro | 0.57290079 |
| 0.01119 | B07 | PE859 | Plate3 | SARS-CoV-2 3CLpro | -4.7963787 |
| 0.08269 | B08 | RO4929097 | Plate3 | SARS-CoV-2 3CLpro | -0.0333082 |
| 0.09519 | B09 | Emricasan | Plate3 | SARS-CoV-2 3CLpro | 0.79939646 |
| 0.07204 | B10 | CGS 27023A | Plate3 | SARS-CoV-2 3CLpro | -0.7427725 |
| 0.08719 | B11 | Talabostat mesylate | Plate3 | SARS-CoV-2 3CLpro | 0.26646549 |
| 0.09014 | C02 | AZVIII-33B | Plate3 | SARS-CoV-2 3CLpro | 0.46298378 |
| 0.07829 | C03 | DMSO | Plate3 | SARS-CoV-2 3CLpro | -0.3264202 |
| 0.07454 | C04 | MDL28170 | Plate3 | SARS-CoV-2 3CLpro | -0.5762316 |
| 0.08274 | C05 | DMSO | Plate3 | SARS-CoV-2 3CLpro | -0.0299774 |
| 0.08389 | C06 | DMSO | Plate3 | SARS-CoV-2 3CLpro | 0.04663146 |
| 0.09039 | C07 | AZVIII-44H | Plate3 | SARS-CoV-2 3CLpro | 0.47963787 |
| 0.08804 | C08 | DMSO | Plate3 | SARS-CoV-2 3CLpro | 0.3230894 |
| 0.08794 | C09 | DMSO | Plate3 | SARS-CoV-2 3CLpro | 0.31642776 |
| 0.08059 | C10 | DMSO | Plate3 | SARS-CoV-2 3CLpro | -0.1732026 |

| | | | | | |
|---|---|---|---|---|---|
| 0.07924 | C11 | DMSO | Plate3 | SARS-CoV-2 3CLpro | -0.2631347 |
| 0.08329 | D02 | DMSO | Plate3 | SARS-CoV-2 3CLpro | 0.00666164 |
| 0.07209 | D03 | Bicailein | Plate3 | SARS-CoV-2 3CLpro | -0.7394417 |
| 0.50684 | D04 | CMPD18-20 | Plate3 | SARS-CoV-2 3CLpro | 28.2220257 |
| 0.23874 | D05 | GC373 | Plate3 | SARS-CoV-2 3CLpro | 10.3621766 |
| 0.07824 | D06 | DMSO | Plate3 | SARS-CoV-2 3CLpro | -0.329751 |
| 0.08409 | D07 | DMSO | Plate3 | SARS-CoV-2 3CLpro | 0.05995473 |
| 0.10644 | D08 | AZVIII-41A | Plate3 | SARS-CoV-2 3CLpro | 1.54883063 |
| 0.44099 | D09 | GC376 | Plate3 | SARS-CoV-2 3CLpro | 23.8353377 |
| 0.09614 | D10 | DMSO | Plate3 | SARS-CoV-2 3CLpro | 0.86268201 |
| 0.04954 | D11 | MWP00709 | Plate3 | SARS-CoV-2 3CLpro | -2.2416409 |
| 0.02289 | E02 | NT 1-32 | Plate3 | SARS-CoV-2 3CLpro | -4.0169672 |
| 0.09329 | E03 | DMSO | Plate3 | SARS-CoV-2 3CLpro | 0.67282535 |
| 0.08949 | E04 | GRL0617 | Plate3 | SARS-CoV-2 3CLpro | 0.41968314 |
| 0.09009 | E05 | DMSO | Plate3 | SARS-CoV-2 3CLpro | 0.45965296 |
| 0.13364 | E06 | AZVIII-34D | Plate3 | SARS-CoV-2 3CLpro | 3.36079593 |
| 0.07969 | E07 | DMSO | Plate3 | SARS-CoV-2 3CLpro | -0.2331573 |
| 0.08989 | E08 | DMSO | Plate3 | SARS-CoV-2 3CLpro | 0.44632969 |
| 0.09419 | E09 | DMSO | Plate3 | SARS-CoV-2 3CLpro | 0.73278008 |
| 0.10229 | E10 | AZVIII-30 | Plate3 | SARS-CoV-2 3CLpro | 1.27237269 |
| 0.08209 | E11 | DMSO | Plate3 | SARS-CoV-2 3CLpro | -0.073278 |
| 0.08309 | F02 | DMSO | Plate3 | SARS-CoV-2 3CLpro | -0.0066616 |
| 0.08234 | F03 | AZVIII-37A | Plate3 | SARS-CoV-2 3CLpro | -0.0566239 |
| 0.09354 | F04 | DMSO | Plate3 | SARS-CoV-2 3CLpro | 0.68947944 |
| 0.07304 | F05 | Betrixaban | Plate3 | SARS-CoV-2 3CLpro | -0.6761562 |
| 0.08944 | F06 | DMSO | Plate3 | SARS-CoV-2 3CLpro | 0.41635232 |
| 0.08344 | F07 | AZVIII-43A | Plate3 | SARS-CoV-2 3CLpro | 0.01665409 |
| 0.00659 | F08 | MAC22272 | Plate3 | SARS-CoV-2 3CLpro | -5.102814 |
| 0.10049 | F09 | Amentoflavone | Plate3 | SARS-CoV-2 3CLpro | 1.15246322 |
| 0.09359 | F10 | DMSO | Plate3 | SARS-CoV-2 3CLpro | 0.69281026 |
| 0.09164 | F11 | DMSO | Plate3 | SARS-CoV-2 3CLpro | 0.56290834 |
| 0.02859 | G02 | Ledipasvir acetone | Plate3 | SARS-CoV-2 3CLpro | -3.6372539 |
| 0.08679 | G03 | Batimastat | Plate3 | SARS-CoV-2 3CLpro | 0.23981894 |
| 0.03034 | G04 | TOFA | Plate3 | SARS-CoV-2 3CLpro | -3.5206752 |
| 0.03934 | G05 | HZ1157 | Plate3 | SARS-CoV-2 3CLpro | -2.9211279 |
| 0.09814 | G06 | Abacavir sulfate | Plate3 | SARS-CoV-2 3CLpro | 0.99591475 |
| 0.09834 | G07 | Sivelestat | Plate3 | SARS-CoV-2 3CLpro | 1.00923803 |
| 0.05434 | G08 | Dasabuvir | Plate3 | SARS-CoV-2 3CLpro | -1.9218823 |
| 0.07869 | G09 | Calycosin | Plate3 | SARS-CoV-2 3CLpro | -0.2997737 |

| | | | | | |
|---|---|---|---|---|---|
| 0.09329 | G10 | 4-Methoxysalicylaldehyde | Plate3 | SARS-CoV-2 3CLpro | 0.67282535 |
| 0.09294 | G11 | Sebacic acid | Plate3 | SARS-CoV-2 3CLpro | 0.64950962 |
| 0.07394 | H02 | Deoxyarbutin | Plate3 | SARS-CoV-2 3CLpro | -0.6162014 |
| 0.10384 | H03 | 2-5-dihydroxyacetophenone | Plate3 | SARS-CoV-2 3CLpro | 1.37562807 |
| 0.09194 | H04 | Oxyresveratrol | Plate3 | SARS-CoV-2 3CLpro | 0.58289325 |
| 0.09649 | H05 | Aloxistatin | Plate3 | SARS-CoV-2 3CLpro | 0.88599774 |
| 0.11434 | H06 | Fostemsavir | Plate3 | SARS-CoV-2 3CLpro | 2.07509997 |
| 0.07149 | H07 | Tasisulam | Plate3 | SARS-CoV-2 3CLpro | -0.7794115 |
| 0.09304 | H08 | Semagacestat | Plate3 | SARS-CoV-2 3CLpro | 0.65617126 |
| 0.07974 | H09 | Triciribine | Plate3 | SARS-CoV-2 3CLpro | -0.2298265 |
| 0.09534 | H10 | IMR-1A | Plate3 | SARS-CoV-2 3CLpro | 0.80938891 |
| 0.09124 | H11 | IMR1 | Plate3 | SARS-CoV-2 3CLpro | 0.53626179 |
| 0.12075 | A02 | Z-IETD-FMK | Plate4 | SARS-CoV-2 3CLpro | 3.10166071 |
| 0.1 | A03 | VR23 | Plate4 | SARS-CoV-2 3CLpro | 1.72277566 |
| 0.08645 | A04 | Amprenavir | Plate4 | SARS-CoV-2 3CLpro | 0.82234711 |
| 0.10545 | A05 | AA26-9 | Plate4 | SARS-CoV-2 3CLpro | 2.08494065 |
| 0.04395 | A06 | Dolutegravir | Plate4 | SARS-CoV-2 3CLpro | -2.0018753 |
| 0.0624 | A07 | Lomibuvir | Plate4 | SARS-CoV-2 3CLpro | -0.7758305 |
| 0.05675 | A08 | Ginsenoside Rh2 | Plate4 | SARS-CoV-2 3CLpro | -1.151286 |
| 0.08095 | A09 | UK371804 | Plate4 | SARS-CoV-2 3CLpro | 0.45685951 |
| 0.02685 | A10 | CA-074 methyl ester | Plate4 | SARS-CoV-2 3CLpro | -3.1382095 |
| 0.0603 | A11 | ML281 | Plate4 | SARS-CoV-2 3CLpro | -0.9153803 |
| 0.1165 | B02 | CP 640186 | Plate4 | SARS-CoV-2 3CLpro | 2.81923847 |
| 0.10795 | B03 | Hydroumbellic acid | Plate4 | SARS-CoV-2 3CLpro | 2.25107138 |
| 0.084 | B04 | Ethyl gallate | Plate4 | SARS-CoV-2 3CLpro | 0.65953899 |
| 0.0811 | B05 | Senegenin | Plate4 | SARS-CoV-2 3CLpro | 0.46682735 |
| 0.0767 | B06 | lithospermic acid | Plate4 | SARS-CoV-2 3CLpro | 0.17443727 |
| 0.0637 | B07 | Dibenzazepine | Plate4 | SARS-CoV-2 3CLpro | -0.6894425 |
| 0.0658 | B08 | LY411575 | Plate4 | SARS-CoV-2 3CLpro | -0.5498927 |
| 0.0668 | B09 | Paritaprevir | Plate4 | SARS-CoV-2 3CLpro | -0.4834404 |
| 0.06765 | B10 | Sofosbuvir | Plate4 | SARS-CoV-2 3CLpro | -0.426956 |
| 0.0935 | B11 | Crenigacestat | Plate4 | SARS-CoV-2 3CLpro | 1.29083576 |
| 0.11605 | C02 | Avagacestat | Plate4 | SARS-CoV-2 3CLpro | 2.78933494 |
| 0.1044 | C03 | Stearic acid | Plate4 | SARS-CoV-2 3CLpro | 2.01516574 |
| 0.09655 | C04 | DMSO | Plate4 | SARS-CoV-2 3CLpro | 1.49351525 |
| 0.5042 | C05 | CMPD18-20 | Plate4 | SARS-CoV-2 3CLpro | 28.5827919 |
| 0.0761 | C06 | DMSO | Plate4 | SARS-CoV-2 3CLpro | 0.13456589 |

| | | | | | |
|---|---|---|---|---|---|
| 0.0685 | C07 | AZVIII-42 | Plate4 | SARS-CoV-2 3CLpro | -0.3704715 |
| 0.0781 | C08 | DMSO | Plate4 | SARS-CoV-2 3CLpro | 0.26747047 |
| 0.07245 | C09 | DMSO | Plate4 | SARS-CoV-2 3CLpro | -0.107985 |
| 0.07745 | C10 | CC42746 | Plate4 | SARS-CoV-2 3CLpro | 0.22427648 |
| 0.07475 | C11 | DMSO | Plate4 | SARS-CoV-2 3CLpro | 0.0448553 |
| 0.07445 | D02 | DMSO | Plate4 | SARS-CoV-2 3CLpro | 0.02491961 |
| 0.07965 | D03 | BTB07789 | Plate4 | SARS-CoV-2 3CLpro | 0.37047153 |
| 0.41755 | D04 | GC376 | Plate4 | SARS-CoV-2 3CLpro | 22.8247009 |
| 0.0346 | D05 | AZVIII-40A | Plate4 | SARS-CoV-2 3CLpro | -2.6232042 |
| 0.04545 | D06 | BTB07420 | Plate4 | SARS-CoV-2 3CLpro | -1.9021968 |
| 0.06985 | D07 | DMSO | Plate4 | SARS-CoV-2 3CLpro | -0.2807609 |
| 0.07115 | D08 | DMSO | Plate4 | SARS-CoV-2 3CLpro | -0.194373 |
| 0.0744 | D09 | AZVIII-44E | Plate4 | SARS-CoV-2 3CLpro | 0.021597 |
| 0.0888 | D10 | DMSO | Plate4 | SARS-CoV-2 3CLpro | 0.97850999 |
| 0.0955 | D11 | MWP00710 | Plate4 | SARS-CoV-2 3CLpro | 1.42374035 |
| 0.1017 | E02 | BTB07421 | Plate4 | SARS-CoV-2 3CLpro | 1.83574456 |
| 0.09715 | E03 | DMSO | Plate4 | SARS-CoV-2 3CLpro | 1.53338663 |
| 0.0795 | E04 | MAC-30731 | Plate4 | SARS-CoV-2 3CLpro | 0.36050368 |
| 0.08255 | E05 | DMSO | Plate4 | SARS-CoV-2 3CLpro | 0.56318317 |
| 0.1028 | E06 | NT 1-24 | Plate4 | SARS-CoV-2 3CLpro | 1.90884208 |
| 0.0716 | E07 | DMSO | Plate4 | SARS-CoV-2 3CLpro | -0.1644694 |
| 0.07325 | E08 | AZVIII-44D | Plate4 | SARS-CoV-2 3CLpro | -0.0548231 |
| 0.0693 | E09 | DMSO | Plate4 | SARS-CoV-2 3CLpro | -0.3173097 |
| 0.07195 | E10 | DMSO | Plate4 | SARS-CoV-2 3CLpro | -0.1412111 |
| 0.0803 | E11 | DMSO | Plate4 | SARS-CoV-2 3CLpro | 0.41366552 |
| 0.0946 | F02 | DMSO | Plate4 | SARS-CoV-2 3CLpro | 1.36393329 |
| 0.09095 | F03 | AZVIII-44B | Plate4 | SARS-CoV-2 3CLpro | 1.12138242 |
| 0.07375 | F04 | DMSO | Plate4 | SARS-CoV-2 3CLpro | -0.021597 |
| 0.0791 | F05 | NT 1-21 | Plate4 | SARS-CoV-2 3CLpro | 0.33392277 |
| 0.0642 | F06 | DMSO | Plate4 | SARS-CoV-2 3CLpro | -0.6562164 |
| 0.076 | F07 | SCR00533 | Plate4 | SARS-CoV-2 3CLpro | 0.12792066 |
| 0.07345 | F08 | DMSO | Plate4 | SARS-CoV-2 3CLpro | -0.0415327 |
| 0.05335 | F09 | Glecaprevir | Plate4 | SARS-CoV-2 3CLpro | -1.3772237 |
| 0.0681 | F10 | SEW03089 | Plate4 | SARS-CoV-2 3CLpro | -0.3970524 |
| 0.07905 | F11 | DMSO | Plate4 | SARS-CoV-2 3CLpro | 0.33060015 |
| 0.0656 | G02 | Doravirine | Plate4 | SARS-CoV-2 3CLpro | -0.5631832 |
| 0.0011 | G03 | Delanzomib | Plate4 | SARS-CoV-2 3CLpro | -4.849356 |
| 0.0675 | G04 | Morroniside | Plate4 | SARS-CoV-2 3CLpro | -0.4369238 |
| 0.04885 | G05 | Calycosin-7-O-beta-D- | Plate4 | SARS-CoV-2 3CLpro | -1.6762591 |

| | | | | | |
|---|---|---|---|---|---|
| | | glucoside | | | |
| 0.05495 | G06 | Glabridin | Plate4 | SARS-CoV-2 3CLpro | -1.2709001 |
| 0.0358 | G07 | Licochalcone A | Plate4 | SARS-CoV-2 3CLpro | -2.5434615 |
| 0.01645 | G08 | Velpatasvir | Plate4 | SARS-CoV-2 3CLpro | -3.8293133 |
| 0.0393 | G09 | Telaprevir | Plate4 | SARS-CoV-2 3CLpro | -2.3108784 |
| 0.0612 | G10 | Odanacatib | Plate4 | SARS-CoV-2 3CLpro | -0.8555733 |
| 0.08295 | G11 | Darunavir | Plate4 | SARS-CoV-2 3CLpro | 0.58976409 |
| 0.0721 | H02 | Danoprevir | Plate4 | SARS-CoV-2 3CLpro | -0.1312433 |
| 0.03045 | H03 | Nelfinavir Mesylate | Plate4 | SARS-CoV-2 3CLpro | -2.8989812 |
| 0.00735 | H04 | Oprozomib | Plate4 | SARS-CoV-2 3CLpro | -4.4340292 |
| 0.0658 | H05 | AEBSF hydrochloride | Plate4 | SARS-CoV-2 3CLpro | -0.5498927 |
| 0.075 | H06 | Belnacasan | Plate4 | SARS-CoV-2 3CLpro | 0.06146837 |
| 0.0761 | H07 | Z-DEVD-FMK | Plate4 | SARS-CoV-2 3CLpro | 0.13456589 |
| 0.1242 | H08 | Z-FA-FMK | Plate4 | SARS-CoV-2 3CLpro | 3.33092112 |
| 0.0698 | H09 | Trovirdine | Plate4 | SARS-CoV-2 3CLpro | -0.2840835 |
| 0.0076 | H10 | MG132 | Plate4 | SARS-CoV-2 3CLpro | -4.4174161 |
| 0.05275 | H11 | Cabotegravir | Plate4 | SARS-CoV-2 3CLpro | -1.4170951 |

**Supplementary Table 3. Structures of synthesized and structurally similar compounds for this study.**

| Compound | Structure |
|---|---|
| AZVIII-30 | 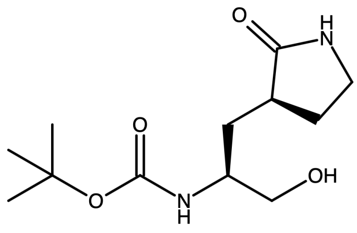 |
| AZVIII-33B | 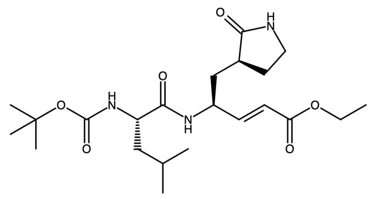 |
| AZVIII-34D | 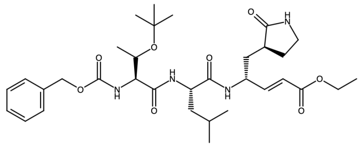 |

| | |
|---|---|
| AZVIII-37A | 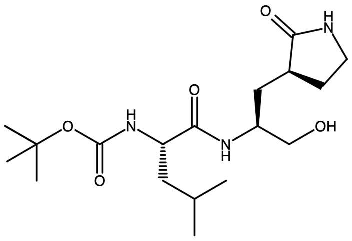 |
| AZVIII-38 | 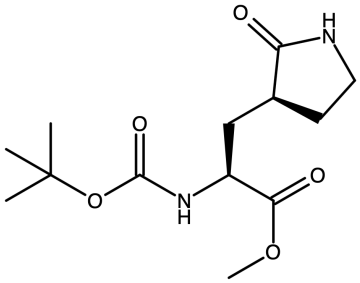 |
| AZVIII-40A | 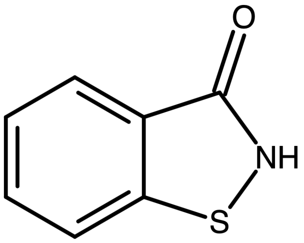 |
| AZVIII-41A | 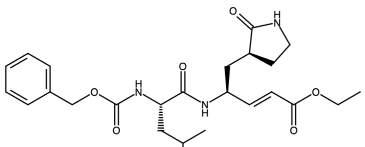 |

| | |
|---|---|
| AZVIII-42 | 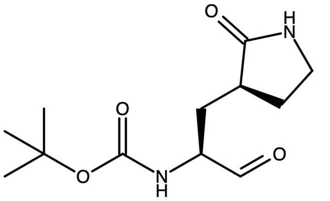 |
| AZVIII-43A | 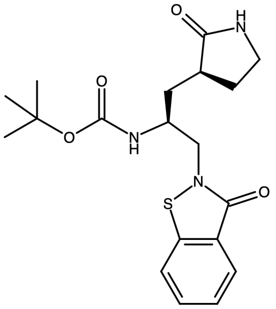 |
| AZVIII-44B | 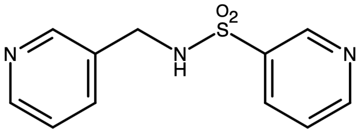 |
| AZVIII-44D | 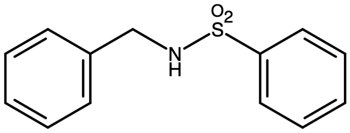 |

| | |
|---|---|
| AZVIII-44E | 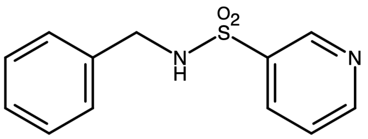 |
| AZVIII-44H | 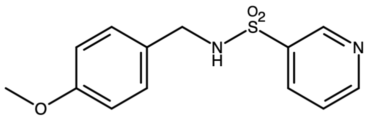 |
| AZVIII-49C | 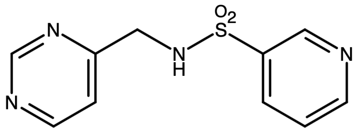 |
| AZVIII-49F | 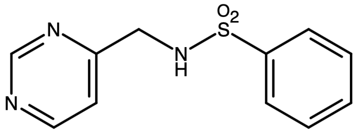 |

| | |
|---|---|
| AZVIII-57D | 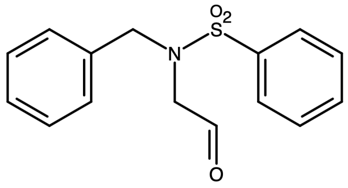 |
| AZVIII-57G | 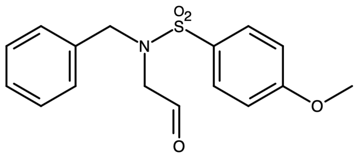 |
| GC373 | 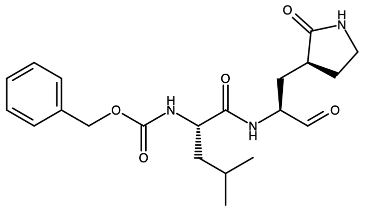 |
| BTB07404 | 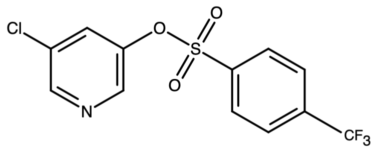 |

| | |
|---|---|
| BTB07407 | 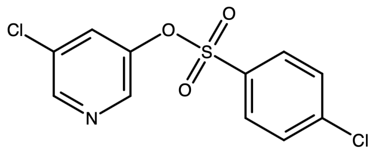 |
| BTB07408 | 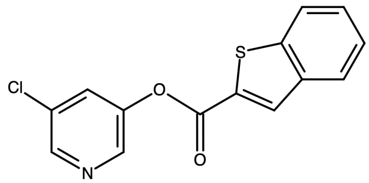 |
| BTB07417 | 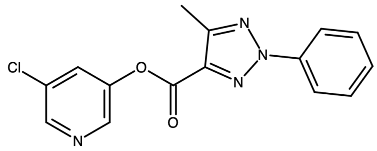 |
| BTB07420 | 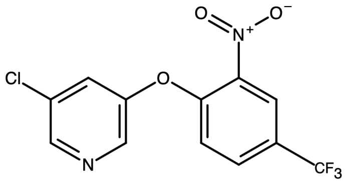 |

| | |
|---|---|
| BTB07421 | 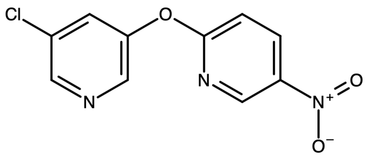 |
| BTB07789 | 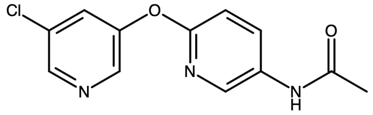 |
| CC42746 | 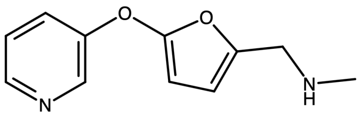 |
| GRL-0496 | 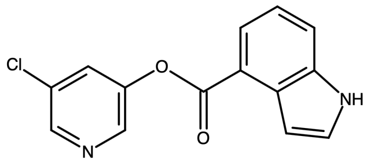 |

| | |
|---|---|
| MAC-30731 | 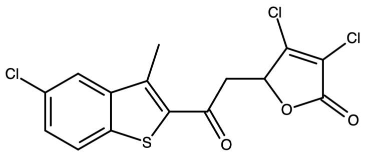 |
| MAC-5576 | 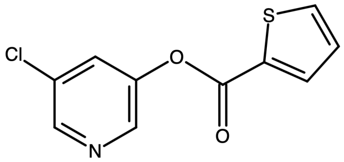 |
| MAC-8120 | 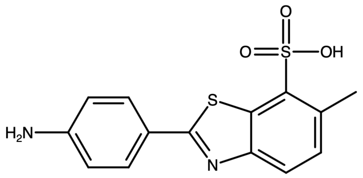 |
| MAC22272 | 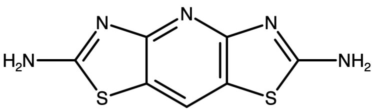 |

| | |
|---|---|
| MWP00332 | 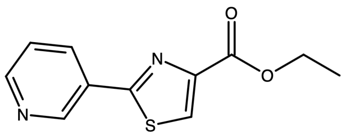 |
| MWP00333 | 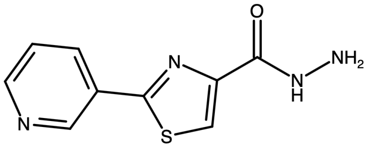 |
| MWP00508 | 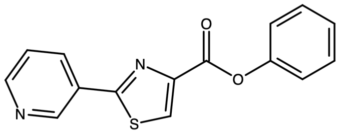 |
| MWP00709 | 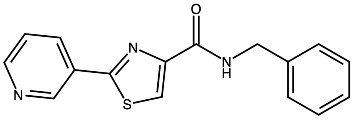 |

| | |
|---|---|
| MWP00710 | 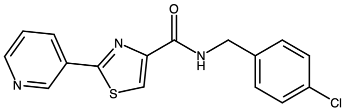 |
| NT 1-21 | 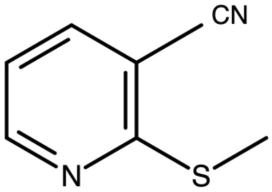 |
| NT 1-24 | 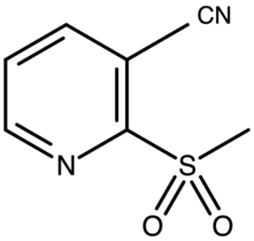 |
| NT 1-32 | 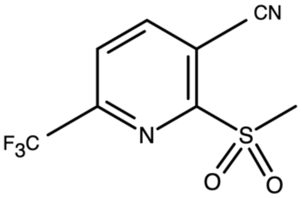 |

| | |
|---|---|
| SCR00533 | 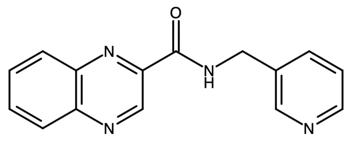 |
| SEW03089 | 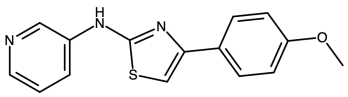 |
| SPB06591 | 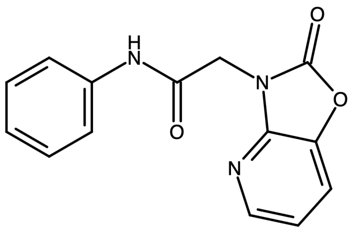 |
| SPB06593 | 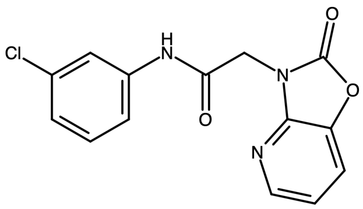 |

| | |
|---|---|
| SPB06613 | 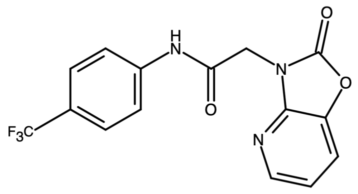 |
| SPB06636 | 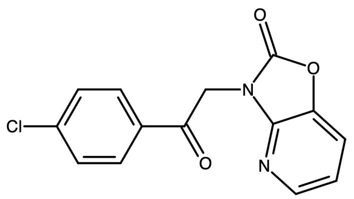 |
| SPB08384 | 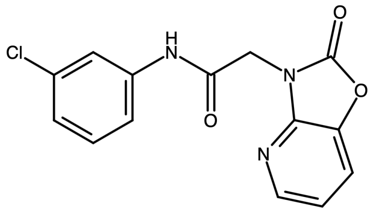 |

**Supplementary Table 4. DNA sequences of proteases used in this study**

| Protease | Sequence |
|---|---|
| SARS-CoV-2 3CLpro | atgagtggttttagaaaaatggcattcccatctggtaaagttgagggttgtatggtacaagtaacttgtggtacaacta<br>cacttaacggtctttggcttgatgacgtagtttactgtccaagacatgtgatctgcacctctgaagacatgcttaaccct<br>aattatgaagatttactcattcgtaagtctaatcataatttcttggtacaggctggtaatgttcaactcagggttattgga<br>cattctatgcaaaattgtgtacttaagcttaaggttgatacagccaatcctaagacacctaagtataagtttgttcgcat<br>tcaaccaggacagactttttcagtgttagcttgttacaatggttcaccatctggtgtttaccaatgtgctatgaggccca<br>atttcactattaagggttcattccttaatggttcatgtggtagtgttggttttaacatagattatgactgtgtctctttttgttac<br>atgcaccatatggaattaccaactggagttcatgctggcacagacttagaaggtaacttttatggaccttttgttgaca<br>ggcaaacagcacaagcagctggtacggacacaactattacagttaatgttttagcttggttgtacgctgctgttataa<br>atggagacaggtggtttctcaatcgatttaccacaactcttaatgactttaaccttgtggctatgaagtacaattatgaa<br>cctctaacacaagaccatgttgacatactaggacctctttctgctcaaactggaattgccgttttagatatgtgtgcttc<br>attaaaagaattactgcaaaatggtatgaatggacgtaccatattgggtagtgctttattagaagatgaatttacacct<br>tttgatgttgttagacaatgctcaggtgttactttccaataa |
| MERS-CoV 3CLpro | atgagcggtttggtgaaaatgtcacatcccagtggagatgttgaggcttgtatggttcaggttacctgcggtagcatg<br>actcttaatggtctttggcttgacaacacagtctggtgcccacgacacgtaatgtgcccggctgaccagttgtctgatc<br>ctaattatgatgccttgttgatttctatgactaatcatagtttcagtgtgcaaaaacacattggcgctccagcaaacttgc<br>gtgttgttggtcatgccatgcaaggcactcttttgaagttgactgtcgatgttgctaaccctagcactccagcctacact<br>tttacaacagtgaaacctggcgcagcatttagtgtgttagcatgctataatggtcgtccgactggtacattcactgttgt<br>aatgcgccctaactacacaattaagggttcctttctgtgtggttcttgtggtagtgttggttacaccaaggagggtagtg<br>tgatcaatttctgttacatgcatcaaatggaacttgctaatggtacacataccggttcagcatttgatggtactatgtatg<br>gtgcctttatggataaacaagtgcaccaagttcagttaacagacaaatactgcagtgttaatgtagtagcttggcttta<br>cgcagcaatacttaatggttgcgcttggtttgtaaaacctaatcgcactagtgttgtttcttttaatgaatgggctcttgcc<br>aaccaattcactgaatttgttggcactcaatccgttgacatgttagctgtcaaaacaggcgttgctattgaacagctgc<br>tttatgcgatccaacaactgtatactgggttccagggaaagcaaatccttggcagtaccatgttggaagatgaattc<br>acacctgaggatgttaatatgcagattatgggtgtggttatgcagtaa |
| SARS-CoV 3CLpro | atgagtggttttaggaaaatggcattcccgtcaggcaaagttgaagggtgcatggtacaagtaacctgtggaacta<br>caactcttaatggattgtggttggatgacacagtatactgtccaagacatgtcatttgcacagcagaagacatgctta<br>atcctaactatgaagatctgctcattcgcaaatccaaccatagctttcttgttcaggctggcaatgttcaacttcgtgtta<br>ttggccattctatgcaaaattgtctgcttaggcttaaagttgatacttctaaccctaagacacccaagtataaatttgtcc<br>gtatccaacctggtcaaacattttcagttctagcatgctacaatggttcaccatctggtgtttatcagtgtgccatgaga<br>cctaatcataccattaaaggttctttccttaatggatcatgtggtagtgttggttttaacattgattatgattgcgtgtctttct<br>gctatatgcatcatatggagcttccaacaggagtacacgctggtactgacttagaaggtaaattctatggtccatttgt<br>tgacagacaaactgcacaggctgcaggtacagacacaaccataacattaaatgttttggcatggctgtatgctgct<br>gttatcaatggtgataggtggtttcttaatagattcaccactactttgaatgactttaaccttgtggcaatgaagtacaac<br>tatgaacctttgacacaagatcatgttgacatattgggacctctttctgctcaaacaggaattgccgtcttagatatgtg<br>tgctgctttgaaagagctgctgcagaatggtatgaatggtcgtactatccttggtagcactattttagaagatgagttta<br>caccatttgatgttgttagacaatgctctggtgttaccttccaataa |
| Bat-CoV-HKU9 3CLpro | atggccggcctgacacgtatggctcacccttcaggtttagtagagccgtgccttgttaaagtaaattatggttccatga<br>ctcttaatggtatatggttggataattttgttatatgtcctaggcatgttatgtgttctagggatgagttagctaatcctgatt<br>accctcgtttgtctatgcgagctgctaattatgattttcacgtgtctcaaaatggtcataatattcgtgttataggccatact<br>atggaaggttcgcttttaaagctaacagttgatgtgaataatcctaaaacacccgcttattcatttatacgggtgagta<br>cgggtcaagctatgagtttgttggcatgttatgatggtttaccaactggtgtgtatacgtgcactttacggtcgaatggt<br>actatgagagcatcattttatgtggctcttgtggtagtcctggctttgtcatgaatggcaaagaagttcaattttgttacct<br>acaccagcttgaattaccaaatggtactcatactggtacagatttttctggtgtcttttatggtccatttgaagacaagc<br>aagtgcctcaattagcggcgcctgattgtactataactgttaatgttttagcatggctttatgcagctgtgttaagtggtg<br>agaattggtttttaaccaagtctagtatttcaccggctgaatttaataattgtgctgttaagtacatgtgtcagtcagtaa<br>cgagtgaaagcttgcaagttttgcaaccacttgcagctaaaacaggtatctctgttgagcgcatgctttcagcgttga<br>aggtattgctctcagctggattttgtggccgtactattatgggctcctgttctttagaggatgagcatacaccgtatgata<br>ttggccgtcaaatgttaggtgttaaattgcaataa |
| HCoV-NL63 3CLpro | atgtctggtcttaagaagatggcacaaccatctggttgtgttgagagatgtgtggttcgcgtctgttatggtagtactgt<br>gcttaatggagtttggttaggtgacactgttacttgtcctagacatgtcatagcaccatcaaccactgttcttattgattat |

| | |
|---|---|
| | gatcatgcatatagtactatgcgtttgcataatttttcagtgtcttataatggtgtcttcttgggagttgttggtgttacaatgcatggttctgtgttgcgtattaaggtttcacaatctaatgtacatacacctaaacatgtttttaaaacgttgaaacctggtgattcttttaatattttagcatgttatgaaggtattgcatctggtgtttttggtgttaatttacgtacaaactttactattaaaggttcttttataaatggagcttgtggttctcctggttataatgttagaaatgatggtactgttgagttttgttatttacaccaaattgagttaggtagtggtgctcatgttggttctgattttactggtagtgtttatggtaattttgatgaccaacctagtttgcaagttgagagtgccaaccttatgctatcagataatgttgttgcctttttgtatgctgctttgttgaatggttgtaggtggtggttgcgttcaactagagttaatgttgatggttttaatgaatgggctatggctaatggttatacaagtgtttctagtgttgagtgctattctattttggcagcaaaaactggtgttagtgttgaacaattgttagcttccattcaacatcttcatgaaggttttggtggtaaaaacatacttggttattctagtttatgtgatgagttcacactagctgaagttgtgaagcagatgtatggtgttaacttgcaataa |
| IBV 3CLpro | atggctggttttaagaaactagttgctcctagtagtgctgttgagaagtgcattgttagtgtctcttatagaggcagtaatcttaatggattgtggttgggtgattccatctactgtccacgacatgtattaggtaagtttagtggtgaccaatggagtgatgtacttagccttgctaataatcatgaatttgaggttgtaactcataatggtgttactttgaatgttgtcagcaggcgtttaaagggtgcattactgattttacagactgcagtagccaatgctgatacaccgaagtataaatttttgaaagcaaattgtggtgatagtttcacaatagcttgctcttatggtggtacagttataggactctaccccgttactatgcgttctaatggaactattagagcatcgtttcttgctggagcatgtggttcagtaggttttaatatagaaaagggtgtagtaaatttttactacatgcaccatctagagttacctaatgcattacacacaggaactgacctaatgggtgagttttatggtggttatatagatgaagaggttgctcagaaagttcaacccgataaattagttactaataatattttggcatggctttatgcagcaattattagtgttaaagagagtagtttttcaacaccaaaatggcttgaaagtactactatcagtattgatgattataataagtgggcaggtgataatggttttacatcatttgtaagctgcactgctattactaaattaagtgctataacaggagtagatgtttgtaaactccttcgtactattatggtaaaaagtgcacaatggggtagtgaccctattttgggacaatataattttgaggatgaaatgacaccggaatctgtttttaatcaggtaggtggtgttaggttacaataa |
| SARS-CoV-2 3CLpro C145A | atgagtggttttagaaaaatggcattcccatctggtaaagttgagggttgtatggtacaagtaacttgtggtacaactacacttaacggtctttggcttgatgacgtagtttactgtccaagacatgtgatctgcacctctgaagacatgcttaaccctaattatgaagatttactcattcgtaagtctaatcataatttcttggtacaggctggtaatgttcaactcagggttattggacattctatgcaaaattgtgtacttaagcttaaggttgatacagccaatcctaagacacctaagtataagtttgttcgcattcaaccaggacagactttttcagtgttagcttgttacaatggttcaccatctggtgtttaccaatgtgctatgaggcccaatttcactattaagggttcattccttaatggttcaGCCggtagtgttggttttaacatagattatgactgtgtctctttttgttacatgcaccatatggaattaccaactggagttcatgctggcacagacttagaaggtaacttttatggaccttttgttgacaggcaaacagcacaagcagctggtacggacacaactattacagttaatgttttagcttggttgtacgctgctgttataaatggagacaggtggtttctcaatcgatttaccacaactcttaatgactttaaccttgtggctatgaagtacaattatgaacctctaacacaagaccatgttgacatactaggacctctttctgctcaaactggaattgccgttttagatatgtgtgcttcattaaaagaattactgcaaaatggtatgaatggacgtaccatattgggtagtgctttattagaagatgaatttacaccttttgatgttgttagacaatgctcaggtgttactttccaataa |
| MERS-CoV 3CLpro C148A | atgagcggtttggtgaaaatgtcacatcccagtggagatgttgaggcttgtatggttcaggttacctgcggtagcatgactcttaatggtctttggcttgacaacacagtctggtgcccacgacacgtaatgtgcccggctgaccagttgtctgatcctaattatgatgccttgttgatttctatgactaatcatagtttcagtgtgcaaaaacacattggcgctccagcaaacttgcgtgttgttggtcatgccatgcaaggcactcttttgaagttgactgtcgatgttgctaaccctagcactccagcctacactttacaacagtgaaacctggcgcagcatttagtgtgttagcatgctataatggtcgtccgactggtacattcactgttgtaatgcgccctaactacacaattaagggttcctttctgtgtggttctGCCggtagtgttggttacaccaaggagggtagtgtgatcaatttctgttacatgcatcaaatggaacttgctaatggtacacataccggttcagcatttgatggtactatgtatggtgcctttatggataaacaagtgcaccaagttcagttaacagacaaatactgcagtgttaatgtagtagcttggctttacgcagcaatacttaatggttgcgcttggtttgtaaaacctaatcgcactagtgttgtttcttttaatgaatgggctcttgccaaccaattcactgaatttgttggcactcaatccgttgacatgttagctgtcaaaacaggcgttgctattgaacagctgctttatgcgatccaacaactgtatactgggttccagggaaagcaaatccttggcagtaccatgttggaagatgaattcacacctgaggatgttaatatgcagattatgggtgtggttatgcagtaa |
| SARS-CoV 3CLpro C145A | atgagtggttttaggaaaatggcattcccgtcaggcaaagttgaagggtgcatggtacaagtaacctgtggaactacaactcttaatggattgtggttggatgacacagtatactgtccaagacatgtcatttgcacagcagaagacatgcttaatcctaactatgaagatctgctcattcgcaaatccaaccatagctttcttgttcaggctggcaatgttcaacttcgtgttattggccattctatgcaaaattgtctgcttaggcttaaagttgatacttctaaccctaagacacccaagtataaatttgtccgtatccaacctggtcaaacattttcagttctagcatgctacaatggttcaccatctggtgtttatcagtgtgccatgagacctaatcataccattaaaggttctttccttaatggatcaGCCggtagtgttggttttaacattgattatgattgcgtgtctttctgctatatgcatcatatggagcttccaacaggagtacacgctggtactgacttagaaggtaaattctatggtccattt |

| | |
|---|---|
| | gttgacagacaaactgcacaggctgcaggtacagacacaaccataacattaaatgttttggcatggctgtatgctg<br>ctgttatcaatggtgataggtggtttcttaatagattcaccactactttgaatgactttaaccttgtggcaatgaagtaca<br>actatgaacctttgacacaagatcatgttgacatattgggacctctttctgctcaaacaggaattgccgtcttagatatg<br>tgtgctgctttgaaagagctgctgcagaatggtatgaatggtcgtactatccttggtagcactattttagaagatgagtt<br>tacaccatttgatgttgttagacaatgctctggtgttaccttccaataa |
| Bat-CoV-HKU9 3CLpro C145A | atggccggcctgacacgtatggctcacccttcaggtttagtagagccgtgccttgttaaagtaaattatggttccatga<br>ctcttaatggtatatggttggataattttgttatatgtcctaggcatgttatgtgttctagggatgagttagctaatcctgatt<br>accctcgtttgtctatgcgagctgctaattatgattttcacgtgtctcaaaatggtcataatattcgtgttataggccatact<br>atggaaggttcgcttttaaagctaacagttgatgtgaataatcctaaaacacccgcttattcatttatacgggtgagta<br>cgggtcaagctatgagtttgttggcatgttatgatggtttaccaactggtgtgtatacgtgcactttacggtcgaatggt<br>actatgagagcatcatttttatgtggctctGCCggtagtcctggctttgtcatgaatggcaaagaagttcaattttgtta<br>cctacaccagcttgaattaccaaatggtactcatactggtacagatttttctggtgtcttttatggtccatttgaagacaa<br>gcaagtgcctcaattagcggcgcctgattgtactataactgttaatgttttagcatggctttatgcagctgtgttaagtgg<br>tgagaattggtttttaaccaagtctagtatttcaccggctgaatttaataattgtgctgttaagtacatgtgtcagtcagta<br>acgagtgaaagcttgcaagttttgcaaccacttgcagctaaaacaggtatctctgttgagcgcatgctttcagcgttg<br>aaggtattgctctcagctggattttgtggccgtactattatgggctcctgttctttagaggatgagcatacaccgtatgat<br>attggccgtcaaatgttaggtgttaaattgcaataa |
| HCoV-NL63 3CLpro C144A | atgtctggtcttaagaagatggcacaaccatctggttgtgttgagagatgtgtggttcgcgtctgttatggtagtactgt<br>gcttaatggagtttggttaggtgacactgttacttgtcctagacatgtcatagcaccatcaaccactgttcttattgattat<br>gatcatgcatatagtactatgcgtttgcataatttttcagtgtcttataatggtgtcttcttgggagttgttggtgttacaatg<br>catggttctgtgttgcgtattaaggtttcacaatctaatgtacatacacctaaacatgtttttaaaacgttgaaacctggt<br>gattcttttaatattttagcatgttatgaaggtattgcatctggtgtttttggtgttaatttacgtacaaactttactattaaagg<br>ttcttttataaatggagctGCCggttctcctggttataatgttagaaatgatggtactgttgagttttgttatttacaccaa<br>attgagttaggtagtggtgctcatgttggttctgattttactggtagtgtttatggtaattttgatgaccaacctagtttgcaa<br>gttgagagtgccaaccttatgctatcagataatgttgttgcctttttgtatgctgctttgttgaatggttgtaggtggtggttg<br>cgttcaactagagttaatgttgatggttttaatgaatgggctatggctaatggttatacaagtgtttctagtgttgagtgct<br>attctattttggcagcaaaaactggtgttagtgttgaacaattgttagcttccattcaacatcttcatgaaggttttggtggt<br>aaaaacatacttggttattctagtttatgtgatgagttcacactagctgaagttgtgaagcagatgtatggtgttaacttg<br>caataa |
| IBV 3CLpro C143A | atggctggttttaagaaactagttgctcctagtagtgctgttgagaagtgcattgttagtgtctcttatagaggcagtaat<br>cttaatggattgtggttgggtgattccatctactgtccacgacatgtattaggtaagtttagtggtgaccaatggagtga<br>tgtacttagccttgctaataatcatgaatttgaggttgtaactcataatggtgttactttgaatgttgtcagcaggcgttta<br>aagggtgcattactgattttacagactgcagtagccaatgctgatacaccgaagtataaattttgaaagcaaattgt<br>ggtgatagtttcacaatagcttgctcttatggtggtacagttataggactctaccccgttactatgcgttctaatggaact<br>attagagcatcgtttcttgctggagcaGCCggttcagtaggttttaatatagaaaagggtgtagtaaatttttactaca<br>tgcaccatctagagttacctaatgcattacacacaggaactgacctaatgggtgagttttatggtggttatatagatga<br>agaggttgctcagaaagttcaacccgataaattagttactaataatattttggcatggctttatgcagcaattattagtg<br>ttaaagagagtagtttttcaacaccaaaatggcttgaaagtactactatcagtattgatgattataataagtgggcag<br>gtgataatggttttacatcatttgtaagctgcactgctattactaaattaagtgctataacaggagtagatgtttgtaaac<br>tccttcgtactattatggtaaaaagtgcacaatggggtagtgaccctattttgggacaatataattttgaggatgaaat<br>gacaccggaatctgtttttaatcaggtaggtggtgttaggttacaataa |
